# Rapid generation of antibody conformational ensembles using machine learning

**DOI:** 10.64898/2026.11.06.764889

**Authors:** Pranav V. Rao, David Sommer, Darcy Davidson, Robert G. Alberstein, Jan Ludwiczak, Joseph Kleinhenz, Bodhi Vani, Eliott Park, Jae Hyeon Lee, Saeed Izadi, James R. Kiefer, Richard Bonneau, Sai Pooja Mahajan, Andrew M. Watkins, Frédéric A. Dreyer

## Abstract

We introduce Cadense, a generative framework for antibody ensemble prediction that produces diverse Fv structures from sequence, informed by conformational dynamics. While a valuable resource, static structure prediction provides only a single snapshot of dynamic proteins, whose motions influence their binding and other properties. This is particularly relevant for antibodies, an important class of molecules for drug discovery, where flexible complementarity determining region (CDR) loops are difficult to model with static approaches. Cadense addresses this gap by generating ensembles that match conformational and biophysical property distributions from molecular dynamics (MD), while reducing the computational burden from days of simulation to seconds of generation. We show Cadense ensembles can recover both *apo* and *holo* conformations of the same antibody and serve as effective input to physics-based docking, capturing the conformational flexibility important to properly model antibody–antigen interactions. Together, these results establish Cadense as a useful tool for antibody conformational ensemble generation with the promise to enable faster, dynamics-aware antibody design.

## 1 Introduction

Antibodies recognize and neutralize pathogens through specific binding to antigens. Therapeutic antibodies exploit these properties: engineered to bind disease-relevant targets with high specificity, they represent the fastest-growing class of biological drugs and have proven crucial in treating cancers, autoimmune disorders, and infectious diseases [1, 2].

The discovery and optimization of therapeutic antibodies has traditionally relied on iterative immunization or display campaigns followed by experimental screening to identify candidates with favorable binding and developability properties [3]. Structure prediction tools, including AlphaFold2 [4] and antibody-specific methods such as ImmuneBuilder [5, 6], IgFold [7], and Ibex [8], have enabled earlier integration of structural information into design workflows [9]. More recently, co-folding models like AlphaFold3 [10], Boltz [11], Chai [12] and Protenix [13] have extended this capability to antibody–antigen complex prediction. The development of these machine learning structure and complex prediction methods has made protein structures more widely available and influential for decision-making in drug discovery. Alongside, structure-based machine learning models have driven progress in *de novo* design [14, 15], allowing for scalable, high-throughput generation of candidate binders against target proteins of interest. Despite this progress, the structural methods underlying these advances operate on static snapshots of inherently flexible, dynamic biomolecules, a limitation particularly consequential for antibodies, where conformational diversity in the CDR loops directly mediates function [16, 17].

Crystallography captures the state most amenable to packing in the lattice, which may or may not be the binding-competent conformation. Blackler et al. illustrated this by solving 20 diverse structures of the same antibody in the absence of antigen. The conformation matching the antigen-bound geometry was already present among these *apo* structures, showing that the *holo* conformation was pre-sampled by antibody-only dynamics [18]. A broader analysis with ITsFlexible shows that approximately 8% of identical CDR sequences crystallize into distinct structural clusters, and MD reveals flexibility in 84% of CDR-H3 loops versus 64% apparent from crystal structures alone [19]. Recent coarse-grained MD simulations further confirm that CDR loops exhibit distinct flexibility profiles often underestimated by static structures [20]. The practical consequences are clear: ensemble-based docking consistently outperforms single-structure approaches [21], and developability-relevant surface properties, such as hydrophobic patches and charge distributions, shift across conformational states in ways that affect candidate ranking [22–24].

While molecular dynamics can indeed capture this conformational diversity, the computational cost of individual simulations makes it impractical for evaluating large candidate libraries [22]. Recently, generative machine learning approaches such as AlphaFlow [25] and BioEmu [26] have begun to address this, moving beyond static outputs to predicting ensembles of protein structures. Boltz-2 [27] also integrates dynamic ensembles in their training pipeline to predict biomolecular complexes. Most directly related to the present work, ABodyBuilder4-STEROIDS generates antibody conformational ensembles with a generative model trained on mostly *coarse grained* antibody molecular dynamics simulations [28].

There are, however, limitations that block direct application of these methods to therapeutic anti-bodies. BioEmu [26] operates on single protein chains, so it cannot model an antibody variable fragment with a paired heavy and light chain. Workarounds that diffuse the heavy chain alone and graft loops onto a static light chain introduce clashes at the interface and discard correlated motions between chains [29]. AlphaFlow [25] shares this same restriction. Both models also rely on multiple sequence alignments (MSAs) for conditioning signal. For antibodies, whose diversity arises from V(D)J recombination and somatic hypermutation rather than evolutionary divergence, an MSA is an unnecessary input to structure-based models, as shown by performant approaches to MSA-free antibody structure prediction [6–8]. Lastly, ABB4-STEROIDS is trained primarily on coarse grained simulations whose coarse grained representation can obscure sidechain and loop dynamics as a tradeoff for scalable simulation [30].

We present Cadense (***C***onformational ***A***ntibody ***D***iffusion ***Ense***mbles), an equivariant diffusion model that generates diverse antibody conformational ensembles directly from sequence, orders of magnitude faster than simulation. Trained on a large dataset of conventional all-atom molecular dynamics trajectories of antibody Fv regions, Cadense samples CDR loop conformations and backbone flexibility. The model operates on generic backbone frames and torsion angles, replacing MSAs with protein language model embeddings [31], and contains no antibody-specific architectural inductive biases beyond the training data. Antibodies, with their flexible loops that influence important binding and biophysical properties [16, 22], serve as a rigorous testbed for validating this MSA-free ensemble generation framework. We evaluate two training variants of Cadense that share the same architecture but differ in their pretraining schemes and how MD trajectories are shown to the model. The primary model, referred to simply as Cadense, uses a curriculum learning strategy where MD frames are weighted by the equilibrium populations of metastable conformational states identified via tICA and PCCA+ clustering of Markov state models (Section 4.4). A second variant, Cadense-*dist*, is finetuned on uniformly sampled MD frames (across time) and is used where reproducing continuous distributional properties of the MD ensemble is the focus.

We demonstrate Cadense’s impact in several ways. First, we show that the generated conformational ensembles span important functional states, recovering both *apo* and *holo* states for antibodies with paired crystal structures, while also capturing the diversity from simulations, e.g., through comparable root mean square fluctuation (RMSF). Next, we show that Cadense-*dist* derived property distributions reduce false-positive risk flags 3.5-fold compared to static structure assessment on our curated benchmark, at a fraction of the computational cost of MD. Lastly we look at physics-based docking with Cadense ensembles, where the model’s ability to flexibly capture diverse CDR conformations leads to significantly improved docking performance over static structures. Together, these results establish Cadense as a practical tool for integrating dynamics into structure-based antibody design.

## 2 Results

Unless otherwise noted, results in this section use the primary Cadense model, trained with the curriculum learning strategy described in Section 4.4, and sampled at normal temperature (see Sec. 4.3) and without guidance. Analyses of per-residue flexibility (RMSF) and biophysical property distributions use Cadense-*dist*, the variant finetuned on uniformly sampled MD frames, which we find better reproduces the continuous distributional characteristics of the simulation ensemble.

### 2.1 Cadense produces realistic conformational ensembles

A key challenge in antibody structure prediction is capturing the conformational flexibility of CDR loops, particularly CDR-H3. Static structure predictors return a single conformation by construction, and while recent diffusion-based models such as AlphaFold3 and Boltz can generate multiple samples, their outputs are typically concentrated around a single dominant state [32]. For antibodies, this state is usually the antigen-bound (*holo*) conformation, reflecting the over-representation of bound structures in training data [33], leaving the *apo* state unrepresented. For applications like docking or property prediction, analyzing both states, and ideally the ensemble connecting them, is crucial [21, 22].

Among the *apo*–*holo* pairs that we held out from training (Sec. 4.5), we identified three pairs with significant conformational changes and single them out to demonstrate that Cadense ensembles span both conformational states (Figure 2). Three cases were held out from Cadense training, while the public PDB structures (`1xf3/2fr4, 4ygv/4yhi, 4s1r`) are present in the training sets of ABodyBuilder3 [6], Boltz-2, and Ibex [8]. We also crystallized the antibody found in *holo* conformation in `4s1r` and arrived at the *apo* structure `Apo 2`, which is unseen by all methods. For each case, we generated 500 Cadense samples (followed by light post-hoc filtering) and compared them to predictions from ABodyBuilder3, Boltz-2, and Ibex (both *apo*- and *holo*-conditioned modes). To compare predicted CDR-H3 loops to both experimental references, we use the angular distance function *d* = 2(1 *−* cos(Δ *θ*)), averaged over backbone dihedrals (*ϕ, ψ*) and loop residues [34, 35]. This alignment-free metric captures local backbone conformation differences; values range from 0 (identical) to 4 (maximally different, i.e. Δ *θ* = *π*) per angle. We plot dihedral distance-to-*holo* against distance-to-*apo* for each predicted structure.

Cadense ensembles consistently produced samples close to both *apo* and *holo* references, spanning the conformational space between them. Baseline methods clustered tightly near the *holo* state, which was included in their training, largely missing the *apo* state. For the Apo 2 *→*4s1r case, where all methods attempt to predict a true out-of-distribution conformation, the best Cadense sample achieved lower dihedral distance to both references than any baseline prediction, and the full ensemble covered the range between states (Figure 2, bottom row). Aggregated across a broader set of seven antibodies with significant conformational changes (ranked by H3 RMSD between *apo* and *holo*), Cadense was the only method to consistently place samples near both references in both dihedral distance and RMSD (Supplementary Figure A1).

A second internally crystallized structure, Apo 1, is likewise unseen by every method evaluated here and shows the same behavior (Figure 1d). The Cadense ensemble recovers the CDR-H3 loop conformation, with the best sample reaching sub-angstrom RMSD to the *apo* state, while Boltz-2, Ibex in *apo* mode and ABodyBuilder3 remain further away in both dihedral distance and RMSD. Together with `Apo 2,` this gives two experimentally determined out-of-distribution cases in which the ensemble contains the crystallographic conformation. In order to provide some labeling to the ensembles, we defined two scores, *centrality* and *support*, described briefly in Fig. 3 and in full detail in Appendix B. As shown in the Figure, restricting to the supported and high centrality subset of the ensemble allowed the identification of structures more similar to the experimental *apo* and *holo* conformations.

**Fig. 1:**
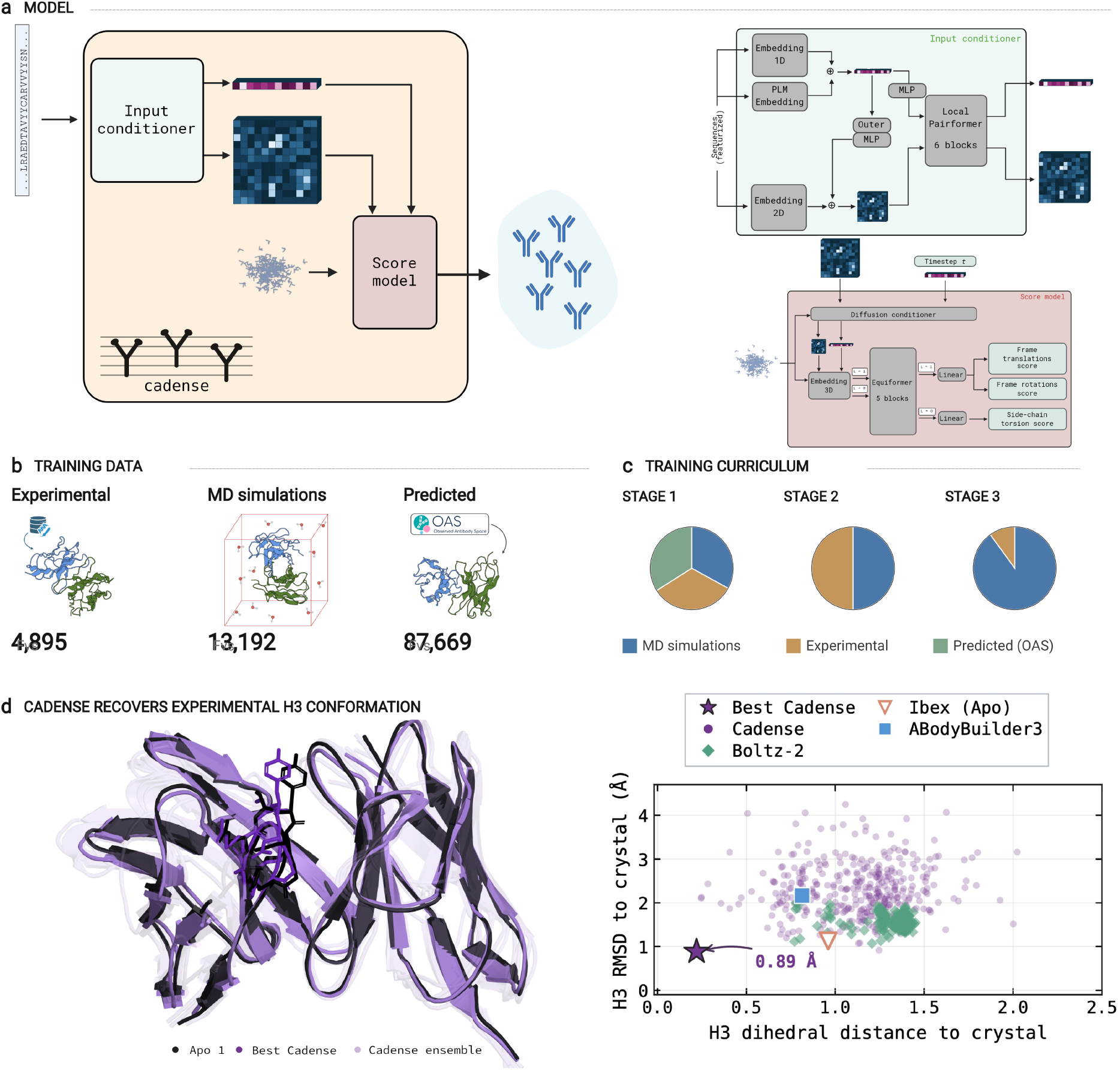
Overview of the Cadense framework for antibody ensemble generation. **(a)** Schematic of the two-component architecture: an input conditioner processes sequence features once, and a score model performs iterative denoising to generate diverse Fv structures. **(b)** Training data sources (experimental structures, MD simulations, predicted structures from pOAS) and **(c)** the three-stage curriculum learning strategy used to train the model. **(d)** Cadense ensemble for an antibody with a newly determined *apo* crystal structure `Apo 1` (PDB 38LM), unseen by all methods. Left: Cadense ensemble (purple) overlaid on the experimental structure (dark blue), showing accurate recovery of the CDR-H3 loop. Right: CDR-H3 dihedral distance versus RMSD to the *apo*-state conformation for Cadense (500 samples), Boltz-2 (500 samples), Ibex (*apo* mode), and ABodyBuilder3; the best Cadense sample (star) achieves sub-angstrom RMSD.

**Fig. 2:**
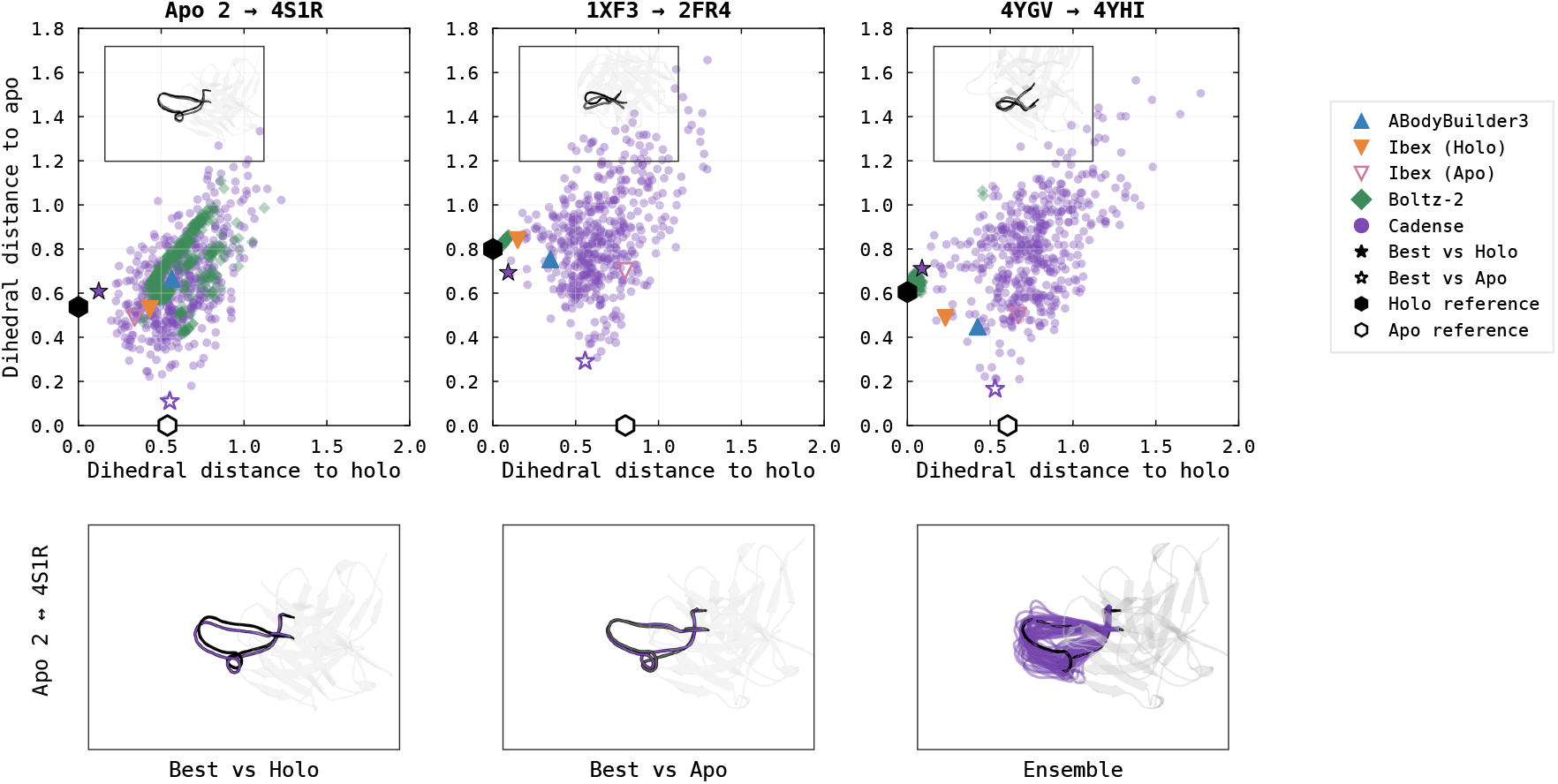
Cadense ensembles capture both *apo* and *holo* CDR-H3 conformations. Top row: dihedral distance of predicted structures to experimental *holo* (*x*) and *apo* (*y*) references for three anti-bodies from our curated set of paired *apo*/*holo* crystal structures. Column titles indicate *apo holo* PDB codes. Cadense samples (*N* = 500) span conformational space between *apo* and *holo* states, while ABody-Builder3, Boltz-2 (*N* = 500) and Ibex produce tighter clusters that may miss one or both states. Stars indicate the closest sample to *holo* (filled) or *apo* (open). Black/white hexagons mark the experimental *holo*/*apo* references. Insets show the CDR-H3 loop of the *holo* reference structure. Bottom row: structural visualization for Apo 2 *→* 4s1r, comparing the best Cadense sample (purple) to the *holo* reference (black, left), *apo* reference (grey, middle), and the full Cadense ensemble (right). Finding the conformation in `Apo 2` (apo state of the `4s1r` antibody; PDB `38LO`) represents an experimental result of the model, as `Apo 2` is a new structure that we crystallized and was therefore unseen by all methods.

**Fig. 3:**
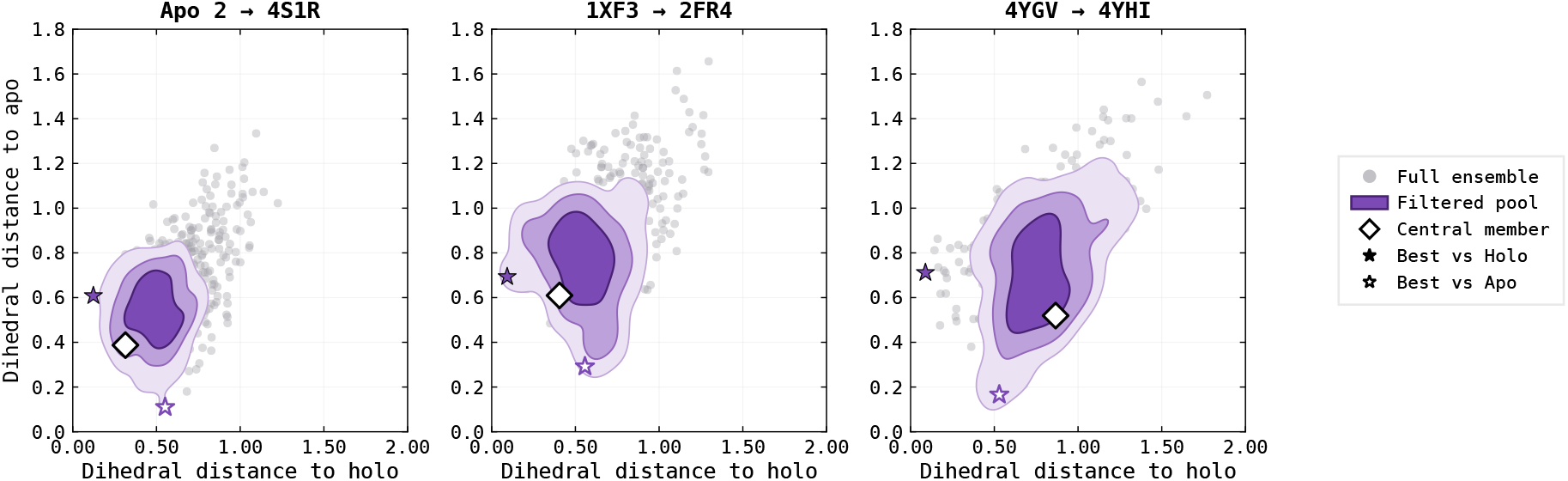
Ensemble-derived scores for locating CDR-H3 conformations closer to experiment. Gray points show the full Cadense ensemble as in Fig. 2. We define two scores, *centrality*, based on dihedral distance to the ensemble medioid, and *support*, a measure of conformational recurrence based on cross-fold nearest-neighbor distances under partitioning of the generated ensemble. The purple distribution summarizes the highest-centrality half of support-passing members, excluding the highest-centrality member shown separately as a white diamond. From light to dark, the contours enclose 95%, 80%, and 50% of the estimated density in this projection.

Beyond capturing discrete *apo* and *holo* states, a useful ensemble model should reproduce the continuous per-residue flexibility profile observed in MD simulations. We evaluated this using Cadense-*dist*, the variant finetuned on uniformly spaced MD frames, by computing the C_*α*_ root-mean-square fluctuation (RMSF) from Cadense-*dist* and Boltz-2 ensembles and correlating it with MD-derived RMSF across 22 held-out antibodies from pOAS, selected to be absent from public structural databases (Figure 4a). Cadense-*dist* ensembles achieved a median Pearson correlation with MD of 0.84, compared to 0.57 for Boltz-2, with notably lower variance. Per-residue traces for representative cases (Figure 4b, c) illustrate that Cadense-*dist* closely tracks the MD flexibility profile across both CDR loops and framework regions, capturing peaks at CDR-H3 and other flexible segments. In contrast, Boltz-2 ensembles substantially underestimate flexibility throughout the Fv, producing near-uniform low RMSF values that fail to distinguish flexible loops from rigid framework. These results confirm that Cadense-*dist* ensembles encode physically meaningful conformational diversity, not merely structural noise, and that this diversity is calibrated to match the dynamics learned from simulation data. In Supplementary Figure A2, we show that Cadense-*dist* ensembles converge to this correlation quickly, within roughly 50 inference structures. We further compare CDR-H3 ensemble diversity against MD for Cadense, Cadense-*dist*, Boltz-2, ABB4-STEROIDS and ITsFlexible in Supplementary Figure A3.

**Fig. 4:**
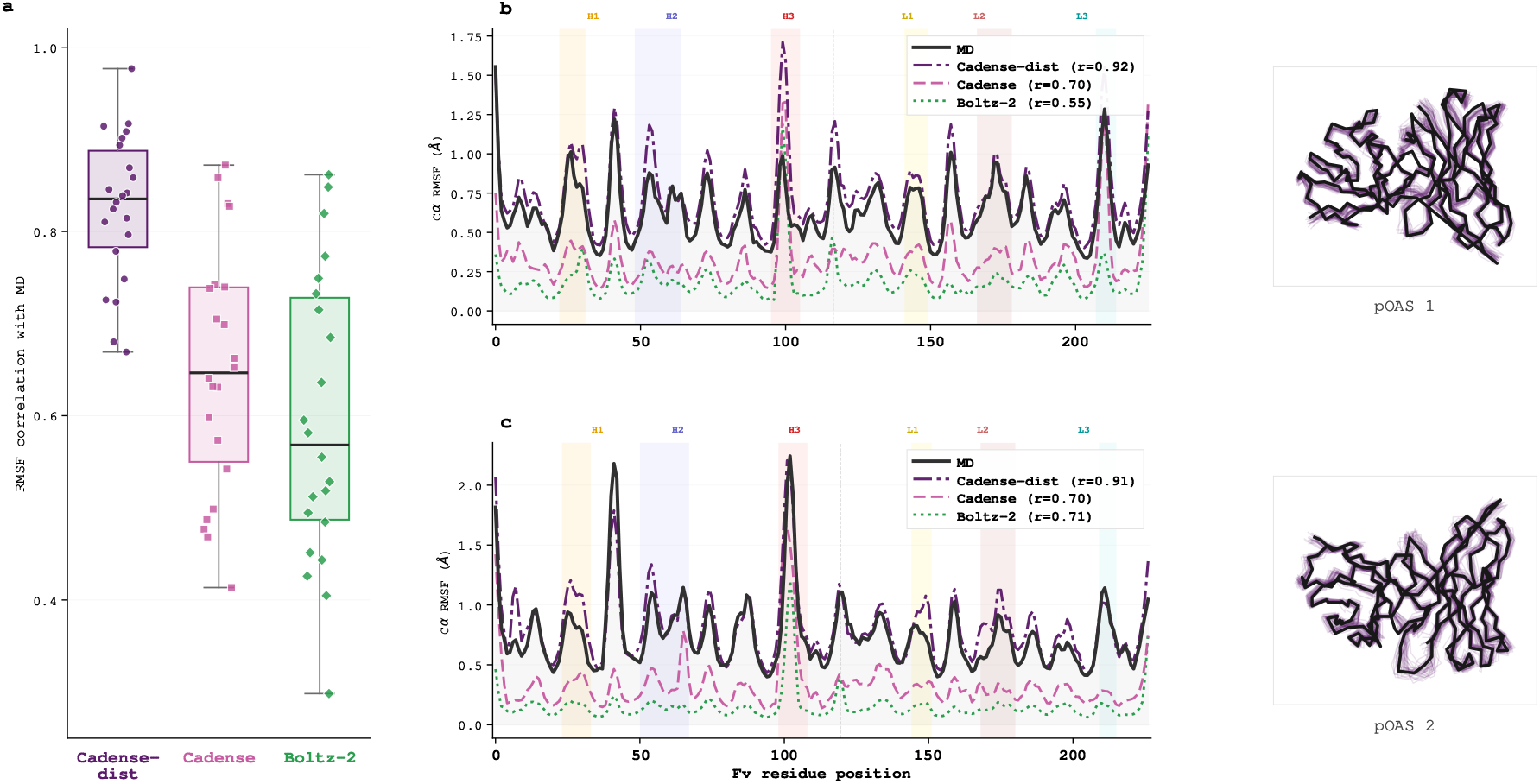
Cadense-*dist* ensembles reproduce per-residue flexibility profiles from molecular dynamics. **(a)** Pearson correlation of per-residue C_*α*_ RMSF between generated ensembles and MD trajectories across 22 held-out pOAS antibodies absent from public structure databases. Cadense-*dist* achieves a median correlation of 0.84 compared to .65 for Cadense and 0.57 for Boltz-2. **(b, c)** Per-residue RMSF for two representative antibodies (`pOAS 1` and `pOAS 2`), comparing MD reference (black), Cadense-*dist* (purple), and Boltz-2 (green). CDR regions are highlighted. Cadense-*dist* tracks the MD flexibility profile across both CDR loops and framework regions, while Boltz-2 substantially underestimates flexibility throughout. Structural insets show the Cadense-*dist* ensemble (purple) overlaid on the MD reference structure (black), illustrating the spatial distribution of conformational diversity.

### 2.2 Cadense ensembles are useful for property flagging and developability

Antibody developability assessment relies on surface properties like hydrophobicity and charge, which are typically computed from static predicted structures. However, these properties vary significantly across conformational ensembles, and static models introduce systematic biases that dynamics can correct [22]. Running MD for every candidate is not tractable at early stages in drug discovery, where screening libraries may contain hundreds to thousands of molecules. We asked whether Cadense-*dist* generated ensembles, which can be sampled in under a minute per antibody (Supplementary Table A1), could replace time consuming MD simulations for property estimation.

We identified *N* = 72 test case antibodies excluded from training. Fifty-five antibodies were chosen from our simulation dataset based on disagreements between ensemble-averaged biophysical properties and those calculated from the first frame of the MD trajectory. The remaining 17 were selected as controls where first-frame and ensemble-averaged properties agreed. Agreement was assessed using risk flags [22]: properties exceeding the 95th percentile of clinical-stage therapeutics are flagged red (high risk), those between the 90th and 95th percentile amber (medium risk), and those below the 90th percentile green (acceptable). A disagreement was defined as any case where the first-frame flag differed from the ensemble-averaged flag for a given property.

Cadense-*dist* ensembles reduced false red flags 3.5-fold compared to static snapshots, here the first frame of the MD trajectory (Figure 5a) on our curated benchmark. Generated ensembles produced only 4 green *→* red misclassifications versus 14 for the first-frame baseline. Overall, Cadense-*dist* ensembles achieved higher accuracy across all flag categories, with particular improvements in green and amber classification and red precision (Figure 5b).

**Fig. 5:**
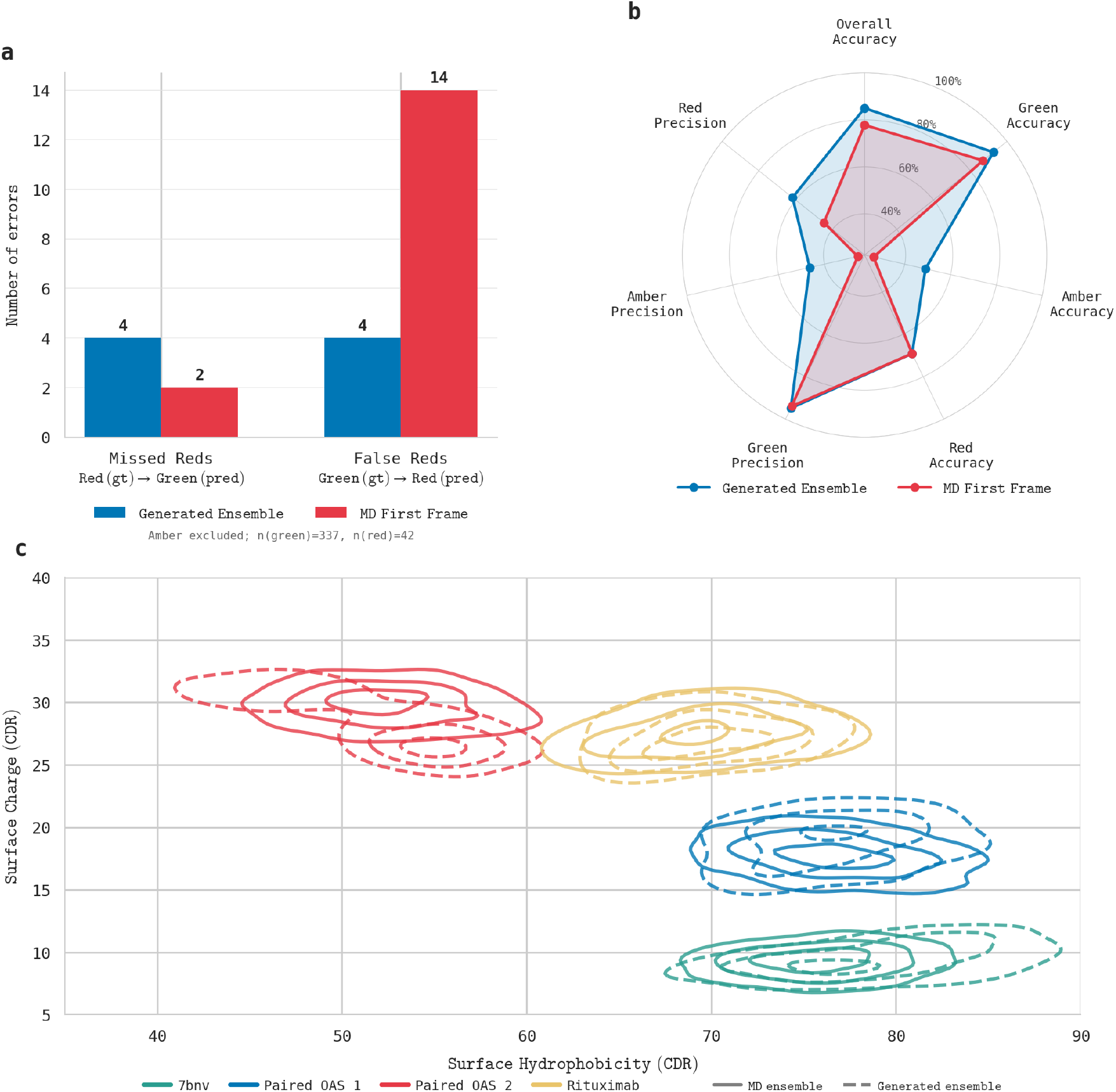
Cadense-*dist* ensembles improve developability risk flag accuracy over single-structure evaluation. **(a)** Critical classification errors compared to the MD ensemble reference. Cadense-*dist* ensembles produce 3.5-fold fewer false red flags (green*→* red misclassifications) than the first-frame baseline, while maintaining comparable missed-red rates. Amber cases excluded; *n* = 337 green, *n* = 42 red reference (MD) flags. **(b)** Radar plot comparing classification performance across flag categories. Cadense-*dist* ensembles (blue) outperform MD first frame (red) in overall accuracy and precision for green, amber, and red classifications. **(c)** Joint distribution of CDR surface hydrophobicity and charge for four representative antibodies spanning diverse regions of property space. Solid contours: MD ensemble; dashed contours: ensemble from Cadense-*dist*. Overlap indicates Cadense-*dist* captures conformational variability relevant to property calculation.

Figure 5c illustrates the distributional agreement between MD and generated ensembles for representative antibodies chosen from the properties test set to cover diverse regions of surface property space, characterized by CDR surface hydrophobicity, computed as a hydrophobic patch score over the CDR surface on the Wimley–White scale (`CDR HPATCH WW`), and CDR electrostatic potential, computed from a

Poisson–Boltzmann treatment (APBS) summed over the CDR surface (`CDR APBS sum`). Contours overlap closely, confirming that Cadense-*dist* ensembles reproduce the property distributions sampled by MD.

### 2.3 Cadense ensembles are effective input to physics-based docking

We now turn to physics-based docking with ensembles, building off the observation that binding-competent paratope conformations can be found in ensembles even starting from *apo* conformations [18, 21], and that this can translate into an effective approach for ensemble-based docking from dynamics.

Here we show that Cadense ensembles capture the conformational diversity relevant to binding, and improve docking outcomes over static predictors. We test this hypothesis at scale on 44 internal antibody– antigen complexes, comparing Cadense ensembles against single-structure baselines ABodyBuilder3 and Ibex-*holo*^1^ and Boltz-2 cofolding, using HADDOCK3 [36] for physics-based docking.

We adapted the ensemble-docking protocol of Ref. [21] to HADDOCK3 to isolate the contribution of paratope conformational sampling from global search. Antibody structures were pre-aligned to the *holo* antigen pose, and the bound antigen conformation was used throughout. To suppress global rigid-body search, we disabled random initial orientations, applied center-of-mass restraints, and limited the rigidbody stage to a single model with reduced van der Waals weighting to prevent rejection of input poses by clash-sensitive scoring prior to refinement. Local interface optimization was performed by flexible refinement (`flexref`) and final energy minimization (`emref`), yielding 50 docked poses per input structure. Ambiguous interaction restraints were defined from the ground truth complex paratope and epitope. For each input structure, we retained the best HADDOCK-scored as that structure’s representative.

For single-structure methods (ABodyBuilder3, Ibex-*holo*) this yields one final prediction. For Cadense, representatives from each ensemble member were then ranked by HADDOCK score, and we evaluated selection at Top-1 and Top-5 (averaging DockQ across the top-ranked poses). We also report an oracle selector that chooses the highest-DockQ representative as an upper bound what a perfect scoring function could extract from the ensemble.

We also include Boltz-2 cofolding as a reference point for full complex prediction. This is purely for a reference noting the difficulty of these models to generalize outside their training distribution [8]. Boltz-2 predicts the complex directly from sequence and is reported here as the best DockQ across 100 seeds.

Cadense ensembles with HADDOCK-based selection outperformed single-structure baselines (Figure 6); broadening selection to the Top-5 docks further improved the score, suggesting that the HAD-DOCK score is a reasonable selection criterion for dock quality. These results indicate that the Cadense ensemble reliably contains high-quality docking solutions, and that simple scoring functions can identify them. However, the oracle performance does offer better performance overall, showing that optimally picking the best complex is still open for further development [38, 39].

**Fig. 6:**
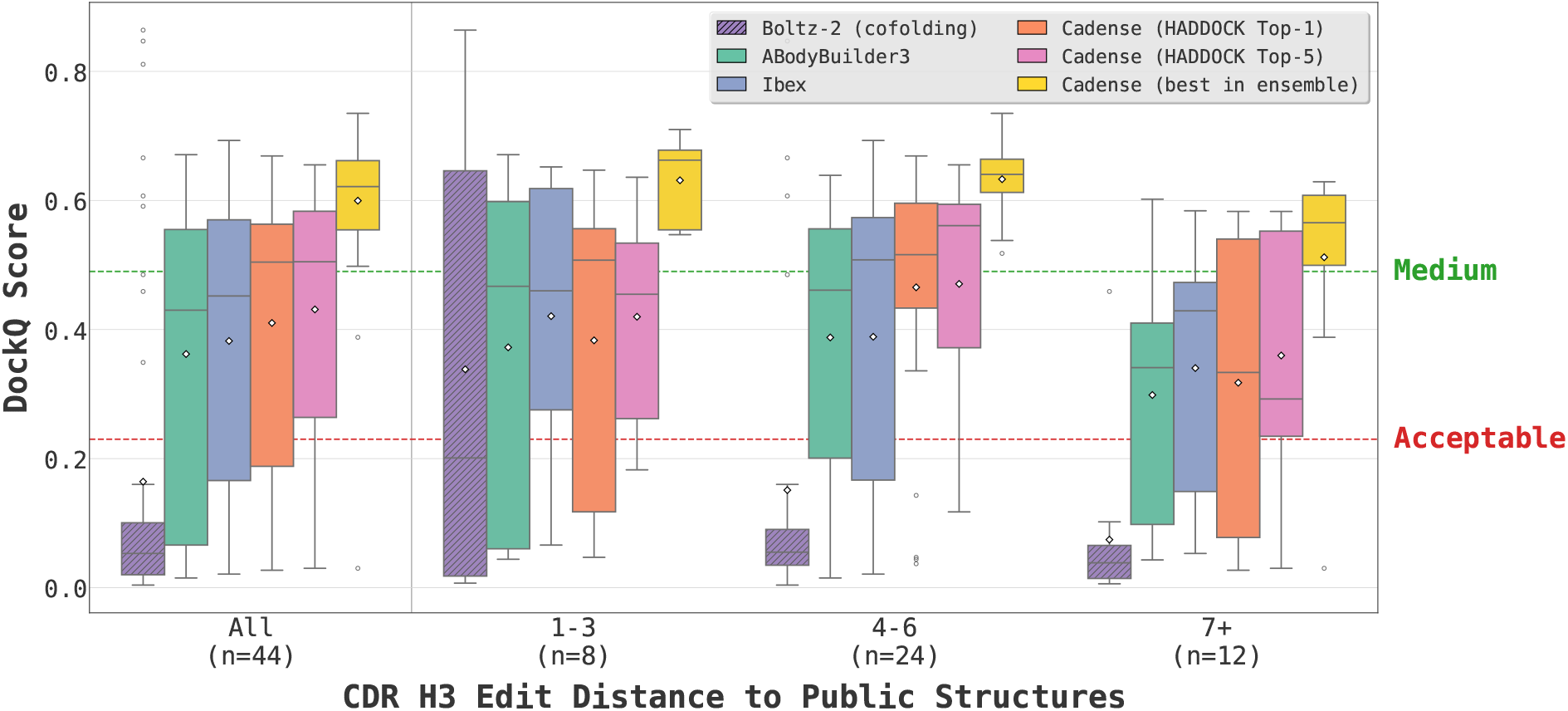
Cadense ensembles improve antibody–antigen docking quality. DockQ scores for *N* = 44 antibody–antigen complexes, shown in aggregate (left) and stratified by CDR-H3 edit distance to the nearest public structure (right). Hatched purple: Boltz-2 cofolding baseline (jointly predicts antibody structure and pose from sequence). Green/blue: single-structure docking predictors (ABodyBuilder3, Ibex). Orange/pink: Cadense ensembles ranked by HADDOCK score, best-scoring pose (Top-1) and mean DockQ across the 5 best-scoring poses (Top-5). Yellow: Cadense oracle (best DockQ in ensemble). Single-structure methods and Boltz-2 degrade on novel CDR-H3 sequences (7+ edit distance), while Cadense ensembles maintain performance. Dashed lines indicate acceptable (0.23) and medium (0.49) quality thresholds [37]. Diamonds indicate mean.

To assess generalization, we stratified results by CDR-H3 edit distance to the nearest public sequence (Figure 6). For antibodies with novel CDR-H3 loops (edit distance 7+), single-structure methods showed degraded performance, while Cadense ensembles maintained higher scores. This suggests that conformational sampling provides robustness to distributional shift, precisely where single-structure predictors struggle most.

## 3 Discussion

Cadense generates diverse conformational ensembles of antibody structures from sequence alone, useful for sampling beyond static structures (including binding-relevant *apo* and *holo* conformations), computing biophysical properties, and as input for docking. While molecular dynamics has demonstrated utility in these areas, Cadense ensembles offer a faster alternative, scalable to early stage antibody discovery and design.

Cadense moves beyond the limitations of existing structure prediction tools, which produce a single conformation by construction or, in the case of generative models trained primarily on crystal structures like Boltz-2, tend to cluster around a dominant mode. Our diffusion ensembles produce structures near both experimental *apo* and *holo* references for held-out test antibodies, most notably for `Apo 1` and `Apo 2`, two new structures we crystallized in-house that were unseen by all methods. We release both to the Protein Data Bank (accession codes `38LM` for `Apo 1` and `38LO` for `Apo 2`) as a resource for benchmarking antibody ensemble generation. Our docking results reinforce this point: diverse sampling, particularly in CDR loops, provides robustness to distributional shift precisely where single-structure methods struggle most, as evidenced by maintained performance on antibodies with novel CDR-H3 sequences (Figure 6). On biophysical properties, Cadense-*dist* demonstrates immediate value. Screening can be a useful tool for antibody discovery and engineering, but false red flags waste resources by eliminating viable candidates at early stages. The 3.5-fold reduction in green *→* red misclassifications translates directly to more calibrated filtering, offering an obvious replacement for computing biophysical properties from static structures without an additional time requirement.

While we provide the centrality and support scores to help label the ensembles, these constructions are still not explicitly labeling which samples may be most *apo*-like or *holo*-like. Learning such a score is a promising direction for future work, potentially through learned selectors trained on docking outcomes [38, 39], which could help recover the oracle-level performance in docking (0.60 against 0.41 mean DockQ). However, even unlabeled, the ensembles have meaningful practical utility. For docking, ranking by HADDOCK score recovers high quality docks from the ensemble candidates, better than static structure baselines. Property estimation operates at the ensemble level, and we show ensemble-averaged flags track MD reference risk flags more closely than any single structure. Lastly, since we demonstrate coverage of the conformational space by recovering experimental conformations (Fig. 2) and correlating with RMSF from dynamics (Fig. 4), this opens up the possibility of flexibility screening for designs. As CDR rigidity is already linked to binding affinity [16, 17], Cadense allows for flexibility to be considered across large libraries of designed antibodies.

Several limitations warrant discussion. Cadense learns the conformational distribution present in its training data and cannot extrapolate to motions or states absent from that data, in particular conformational changes on timescales beyond those sampled with our short 200 ns conventional simulations. Accordingly the model also does not yield kinetic information, which is another extension we leave for future work. Our docking benchmark uses *holo* antigen structures for pre-alignment [21] and restraints for both paratope and epitope, so performance in more general settings remains to be considered. Finally, the current model operates on Fv regions only; extension to full-length antibodies or other modalities would require an expanded simulation dataset for training. In the same vein, the current framework models antibody dynamics in isolation; explicitly incorporating the antigen and modeling conformational dynamics on the target side is a natural extension, though it would likewise require corresponding complex simulation data.

Looking forward, incorporating experimental observables from nuclear magnetic resonance [40] or hydrogen–deuterium exchange [41] for inference-time guidance [42] or train-time experimental finetuning [26] could aid the diversity and generalizability of generated ensembles. More broadly, this work suggests that generative models trained on simulation data can serve as efficient surrogates for physics-based sampling across protein classes where conformational flexibility impacts function and throughput constraints limit MD usage. Cadense enables routine access to antibody conformational ensembles, allowing for important aspects of the dynamic nature of antibodies to inform earlier stages of antibody design and discovery.

## 4 Cadense model

In this section we describe the model architecture, diffusion formulation, sampling procedure, and training data used to build Cadense; full algorithmic details and derivations are provided in the supplementary materials.

### 4.1 Architecture overview

Cadense follows a two-component design (Figure 1a): (1) an *input conditioner* that performs initial featurization through compute-intensive layers, and (2) a lighter-weight *score model* that performs denoising diffusion of protein structure. This split amortizes the cost of the input conditioner, which is evaluated once per sequence, across all diffusion timesteps.

We represent protein geometry using backbone rigid frames for each residue *i*—translations *t*_*i*_ *∈* ℝ^3^ and rotations *r*_*i*_ *∈* SO(3) derived from C_*α*_, N, and C atoms ala AlphaFold2 [4]—together with five torsion angles *χ*_*iq*_ *∈* SO(2), *q ∈ {*0, …, 4*}*, per residue (one backbone oxygen torsion and four side-chain dihedrals).

The input conditioner initializes single (per-residue) and pair (per-residue-pair) representations from sequence features, including protein language model embeddings [31] and relative position encodings, then refines them through Pairformer-style attention blocks analogous to AlphaFold2 and AlphaFold3 [4, 10]. To improve computational efficiency, we use a form of semi-local triangle attention and triangle multiplication operations inspired by Proteus [43], reducing the naїve *O* (*n*^3^) complexity to *O* (*kn*^2^) for coordination number *k ≤ n* and total number of residues *n*.

The score model is built on the Equiformer architecture [44], a graph attention network incorporating SE(3)-equivariant components based on irreducible representations of SO(3). We restrict steerable features to maximum order *L*_max_ = 1 (scalars and vectors) and use depth-wise Clebsch–Gordan tensor products to manage computational cost in the interaction layers. The message-passing graph is constructed using *k*-nearest neighbors supplemented with random long-range edges sampled with inverse-cubic radial propensity, following Chroma [45]. Following Equifold [46], we employ a hierarchical message-passing scheme with three node types: backbone nodes (one per residue), side-chain nodes (each connected only to their parent backbone node), and a global auxiliary node fully connected to all backbone nodes that maintains a latent representation of the system state (Appendix F).

### 4.2 Diffusion formulation

Structure generation is realized by a joint Riemannian diffusion model over the product manifold SE(3) *×* SO(2)^5^ defined by rigid backbone frames, SE(3) = SO(3) ⋉ ℝ^3^, and side-chain torsion angles [47]. For backbone translations, we use a variance-preserving (VP) schedule with a geometric progression [48], constraining translations to the zero center-of-mass subspace [49]. The stationary distribution is a zero-center-of-mass projected normal distribution with isotropic covariance scaled by physical length units. For backbone rotations, we use a variance-exploding (VE) schedule [50] where the forward process corresponds to Brownian motion on SO(3), and the stationary distribution is the Haar measure, i.e., the high-noise limit of the IGSO(3) heat kernel [49]. Side-chain torsions similarly follow a VE schedule on SO(2), where the stationary distribution is again the Haar measure, in this case the high-noise limit of the wrapped normal distribution [51].

### 4.3 Sampling

#### Reverse diffusion

Baseline sampling uses Euler–Maruyama discretization of the time-reversed SDEs over SE(3) *×* SO(2)^5^. Following Chroma [45], we implement hybrid Langevin dynamics to control the trade-off between sample diversity and quality. This introduces a temperature-like rescaling parameter *λ*_0_, where *λ*_0_ *>* 1 emphasizes high-likelihood states, together with an equilibration factor *ψ >* 0 that introduces annealed Langevin dynamics to improve mixing across multimodal distributions. When guidance potentials are applied (see below), we include up to 5 corrector steps per predictor step following annealed Langevin dynamics [50] to ensure adequate mixing under the additional gradient fields.

#### Guidance

At generation time, auxiliary guidance potentials 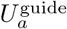 can be composed with the learned score to steer sampling toward desired structural properties. Gradients of these potentials are incorporated into both predictor and corrector steps of the reverse dynamics. We use guidance based on AlphaFold2 structure violation losses [4] to reduce steric clashes in generated structures; further details and the impact of guidance on sample quality are provided in Appendix H.3.

#### Post-hoc quality control

Generated structures pass a post-hoc geometric quality filter (Appendix A.1); comparator methods are evaluated unfiltered, and rejected samples count against the Cadense generation budget.

#### Sampling configuration in results

Unless otherwise noted, results use the normal-temperature setting *λ*_0_ = 1, *ψ* = 0 without guidance. Generated structures pass a post-hoc geometric quality filter (Appendix A.1); comparator methods are evaluated unfiltered, and rejected samples count against the Cadense generation budget.

### 4.4 Training data and strategy

Cadense is trained on molecular dynamics trajectories of 13,192 unique antibody Fv sequences. Sequences were drawn from SAbDab [52], TheraSAbDab [53], and pOAS [54], with subselection strategies designed to maximize diversity in sequence space and surface properties (Appendix D). Initial structures were predicted with ESMFold [31] and refined with Rosetta [55], then used to initiate 200 ns molecular dynamics simulations using the Amber FF14SB force field [56] with TIP3P water (Appendix D).

For training, we extract frames from the production trajectories and represent each as backbone frames and torsion angles. We evaluate two strategies for frame selection during the MD finetuning stage. The primary model (Cadense) uses a curriculum learning strategy, and trajectories are clustered before training. For clustering, we first projected CDR backbone dihedral features with tICA [57], constructed a Markov state model, and then applied PCCA+ [58] clustering to identify metastable conformational states; all of this was done using the PyEMMA package [59] (Appendix E). Training frames are then sampled proportionally to the stationary distribution of the MSM, weighting the data toward equilibrium-populated metastable states. The second variant (Cadense-*dist*) uses uniformly spaced frames from each trajectory, providing broader temporal coverage of the simulation ensemble. Both variants share the same architecture and have one stage of crystal-structure pretraining. In contrast to the three-stage curriculum strategy of the base Cadense model, Cadense-*dist* has a pretraining stage on crystal structures (no synthetic data) and is finetuned on equally spaced MD frames. We find the first recovers discrete functional states more sharply, while the second better reproduces continuous distributional quantities such as RMSF and biophysical property distributions.

- **Cadense**: base model, three-stage curriculum learning, uses clustered MD frames for training
- **Cadense-*dist*** : MD distribution specialist model, pretrained on experimental crystal structures, finetuned on equally spaced MD frames

Both models are trained to denoise structures corrupted according to the noise schedules described in Appendix G.1, with a piecewise-linear timestep sampling scheme that samples uniformly for *t >* 0.25 (below which structures are essentially resolved given the noise schedules).

### 4.5 Held-out evaluation sets

Three held-out sets were constructed for the analyses in Section 2:

1. **Apo/holo pairs**. A set of 33 antibodies with paired *apo* and *holo* crystal structures.
2. **Biophysical properties and RMSF evaluation**. A set of 72 antibodies chosen for biophysical property analysis. The selection criteria are further detailed in Section 2.2; a subset of these structures (N=22 antibodies from pOAS) was used for the RMSF evaluation in Fig 4.
3. **Internal crystal structures**. Two Fab structures, Apo 1 and Apo 2, expressed, purified and crystallized in-house (Appendix C).

To prevent data leakage between training and our evaluation sets, we clustered all the sequences from the curriculum learning dataset in Fig. 1 against the held-out sequence sets using MMSeqs2 [60] at 80% sequence identity, and clusters overlapping with the test set were also held out from training.

## Acknowledgements

We thank the staff of the Shanghai Synchrotron Radiation Facility (SSRF) beamline BL02U1, where diffraction data were collected and Yanshu Dou and colleagues at Viva Biotech for work related to the crystal structures. We thank Feng Wang and Hong Li for their coordination of protein purifications.

## Appendix A Supplementary tables & figures

**Fig. A1:**
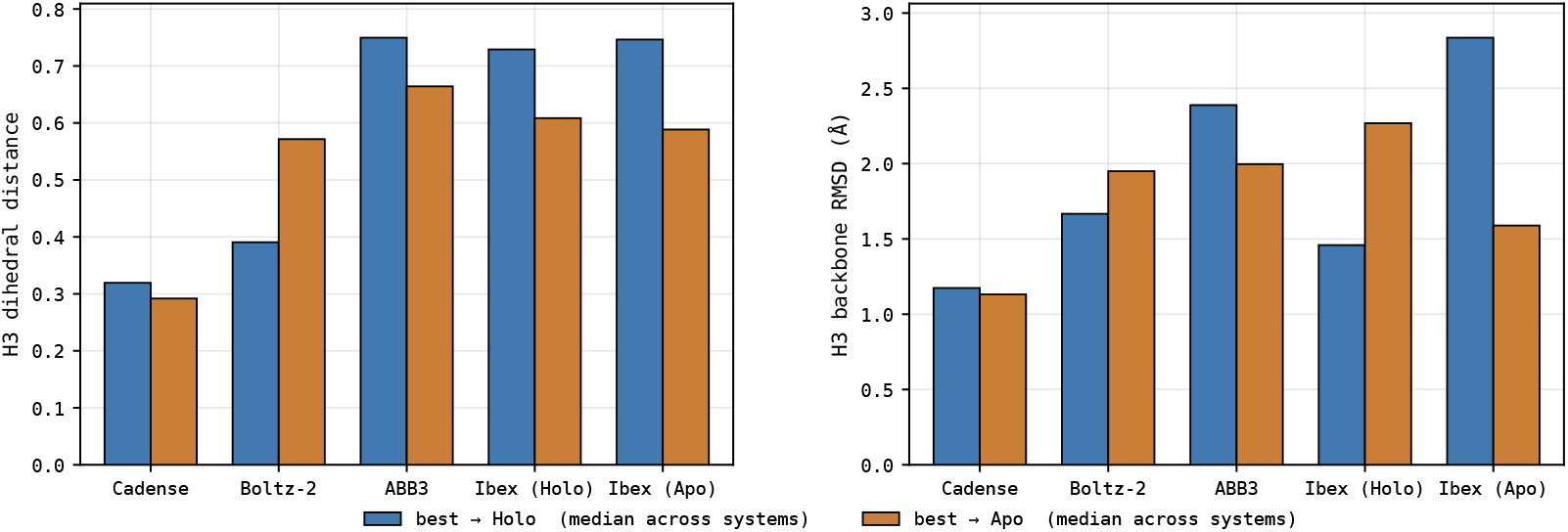
Aggregated apo/holo coverage across seven medium-hard apo/holo examples. We ranked curated antibodies with resolved apo and holo structures by H3 RMSD and chose seven with the largest H3 RMSDs, a superset of those compared in the three highlight examples in Fig. 2. For each method, we report the smallest distance among its included predictions to the *holo* reference (blue) and, independently, to the *apo* reference (orange). We select the best-to-holo and best-to-apo samples independently, so the two samples generally differ for the ensemble methods (Cadense: 500 samples/system; Boltz-2: 500 samples/system). ABB3 and each Ibex variant contribute one prediction per system. **Left:** H3 dihedral distance. **Right:** H3 backbone RMSD (Å), calculated over N, C_*α*_, C, and O atoms after superposition on the heavy-chain framework backbone. We report the median across the seven systems; lower values indicate closer agreement with the corresponding experimental state. The text below the panels gives the median experimental apo*↔* holo reference separation under each metric: 0.54 for dihedral distance and 3.27 Å for H3 backbone RMSD. At the matched 500-sample budget, Cadense has the lowest median best-to-reference distance for both endpoints under both metrics.

### A.1 Structural quality control of generated antibody ensembles

We applied post-generation structural quality control (QC) to Cadense-generated Fv structures. Input structures were standardized to contain heavy and light chains labeled H and L. A structure was retained only if it passed all of the following checks:

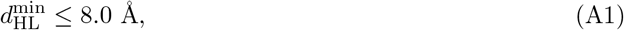

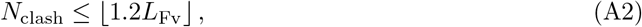

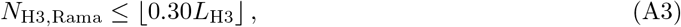

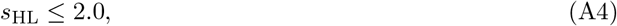

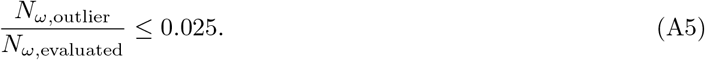

Here, 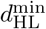 is the minimum inter-chain C_*α*_–C_*α*_ distance; *L*_Fv_ is the number of heavy- and light-chain C_*α*_ residues; and *N*_clash_ is the number of atom pairs belonging to different residues that are separated by at most 2.0 Å. The clash criterion (second line) therefore sets a clash budget that scales with system size, retaining a structure only if it contains at most 1.2 such clashing atom pairs per Fv residue; it limits steric overlaps and is not a filter on the number of residues.

CDR-H3 residues were defined using AHo numbering. For each evaluable H3 residue, the backbone *ϕ* and *ψ* angles were compared with the 90 *×* 90 MDAnalysis Ramachandran reference grid. Residues outside the retained reference contour (grid value below 1.0, corresponding to the MDAnalysis 99% reference contour) were counted as Ramachandran outliers. The permitted number of outliers therefore scaled with H3 length, *L*_H3_.

**Fig. A2:**
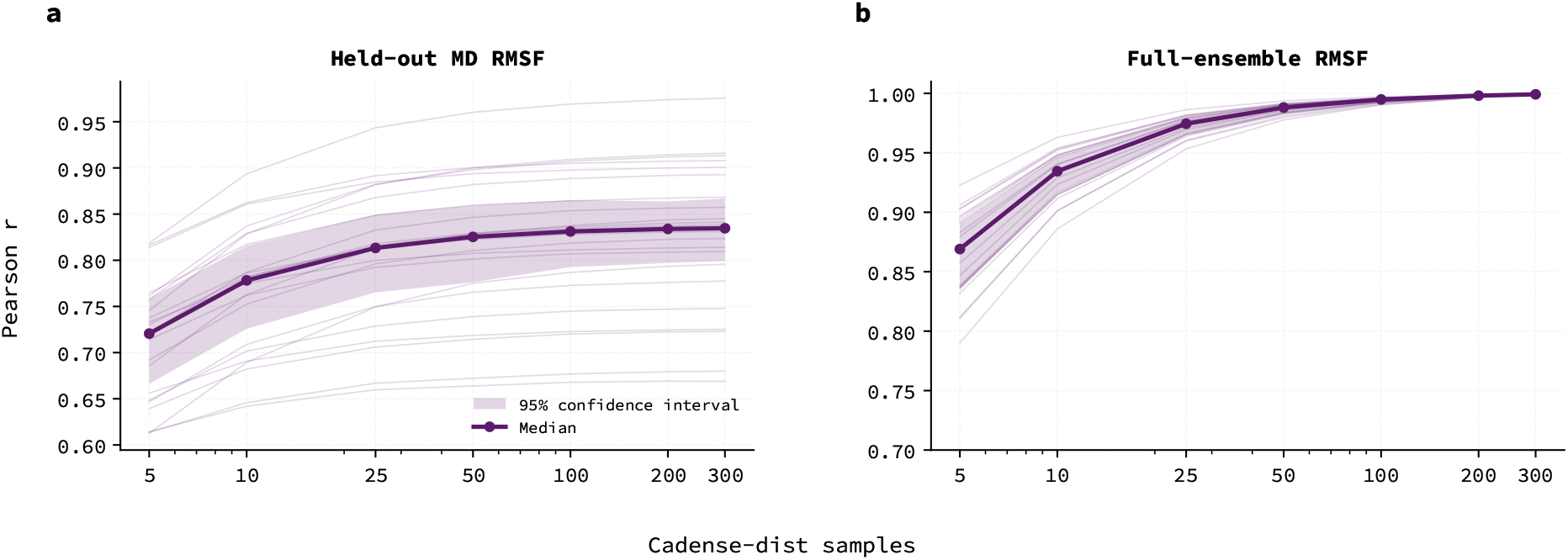
Ensemble property convergence through sampling: In this figure we show how ensemble properties stabilize relatively quickly through sampling from the model, shown by considering RMSF computed from Cadense-*dist* ensembles of increasing size. In **a)** we show that the median RMSF correlation across the N=22 set of antibodies from pOAS settles with relatively small Cadense-*dist* ensemble size. Faded lines represent RMSFs for individual pOAS antibodies versus ensemble size. In **b)** we compare the truncated Cadense-*dist* ensembles to the full ensemble for each antibody, depicting the same convergence phenomenon.

**Table A1:** Runtime comparison. Wall-clock comparison for Apo 1. Cadense and Boltz-2 each use 100 raw generation attempts on one NVIDIA B200; explicit GPU warmup is excluded. Cadense input time is target conditioning, and post-hoc filtering is applied after generation using eight CPU workers. Boltz-2 input time is the measured local MSA search, added once per target. The MD row is the documented 200-ns production trajectory (approximately 12 h on one A100), not 100 independent MD samples; MD system preparation, equilibration, frame selection, and post-simulation quality control were not timed. Consequently, the MD speedups are not hardware-normalized and should be interpreted as workflow-level wall-time comparisons.

| Method | Output | Input / conditioning | Generation / simulation | Updated QC | Total | Speedup vs. MD |
|---|---|---|---|---|---|---|
|  |  | Stage wall time (s) |  |  |  |  |
| Cadense | 100 raw<br>66 retained | 4.3 | 412.3 | 0.8 | 417.3 | 103.5× |
| Boltz-2 | 100 raw | 884.0 | 124.7 | — | 1,008.7 | 42.8× |
| MD | one 200-ns trajectory | — | 43,200.0 | — | 43,200.0 | 1.0× |

The dimensionless VH–VL separation score was defined as

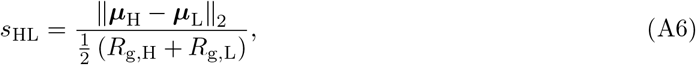

where ***µ***_H_ and ***µ***_L_ are the C_*α*_ centroids and *R*_g,H_ and *R*_g,L_ are the corresponding C_*α*_ radii of gyration. Peptide-bond planarity was evaluated over both chains using *ω* = C_*α,i*_–C_*i*_–N_*i*+1_– C_*α,i*+1_. A peptide bond was classified as an omega outlier when its dihedral was more than 30^*°*^ from both the cis (0^*°*^) and trans (*±*180^*°*^) states. Structures with no evaluable omega angles, or with any parsing or scoring error, failed QC. All thresholds were inclusive.

QC was treated as part of the Cadense generation protocol and was therefore used to filter only Cadense structures. Comparator methods were evaluated without post-hoc structural filtering. Cadense structures rejected by QC still counted toward the raw generation budget, and endpoint statistics were calculated from the retained Cadense structures.

**Fig. A3:**
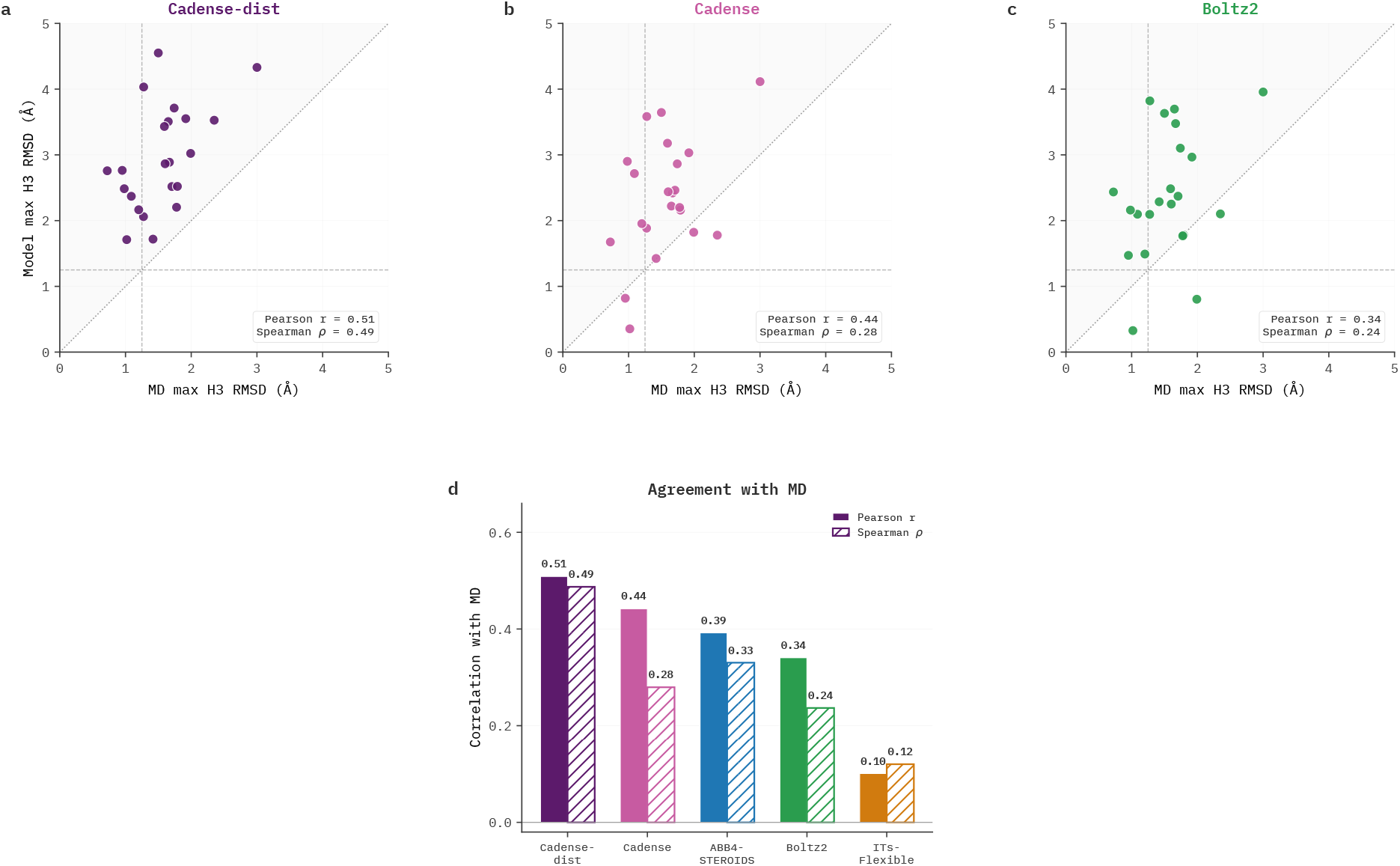
Comparison of generated CDR-H3 ensemble diversity with molecular dynamics. (a–c) We evaluated 22 held-out pOAS antibodies by plotting the exact maximum pairwise C_*α*_ RMSD of each generated CDR-H3 ensemble against the same statistic from the MD trajectory. For Cadense-*dist*, Cadense, Boltz-2, and ABB4-STEROIDS [28]we generated 500 structures per target (QC was applied to Cadense ensembles). In the first row, each dot represents one antibody structure, and the dotted diagonal marks equality; the horizontal and vertical dashed lines mark the 1.25 Å threshold that ITsFlexible [19] uses to distinguish CDR conformational states. Insets report Pearson and Spearman correlations across targets. **(d)** Bars report the Pearson correlation between each method’s flexibility measure and the MD-derived maximum CDR-H3 RMSD. We ran ITsFlexible on one ABodyBuilder2-predicted structure per target and used its scalar flexibility score because ITsFlexible does not generate an ensemble.

## Appendix B Ensemble-derived centrality and support scores

We use *centrality* to describe a generated member’s position relative to the center of its ensemble and *support* to describe the recurrence of similar conformations across subsets of that ensemble. We calculate both scores within each antibody’s QC-passing ensemble using CDR-H3 *ϕ, ψ* angles. The calculation uses generated structures, sequence-derived residue correspondence, and sample identifiers; experimental coordinates and distances to the apo/holo references enter only the subsequent visualization and evaluation.

### B.1 Centrality (closeness to ensemble medoid)

For an ensemble of *n* members, let *θ*_*it*_ denote angle *t* of member *i*, with *T* = 2*L* angles for a CDR-H3 loop of *L* residues. We match CDR-H3 residues using AHo numbering and calculate the pairwise (square-root) dihedral distance

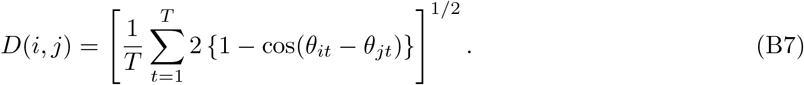

This distance represents each angle by its sine and cosine and respects angular periodicity; we use this same metric to compare ensemble members distance to experimental *apo* and *holo* conformations in Fig. 2.

We first identify the ensemble medoid, the generated member with the smallest mean distance to the other members:

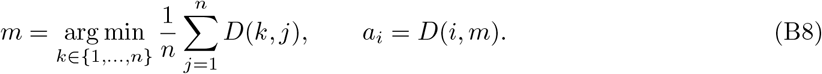

We then assign centrality by the empirical rank of *a*_*i*_, with higher scores for members closer to the medoid:

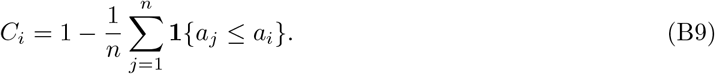

Here **1** {·} is the indicator function. The ranking includes the member itself and assigns equal centrality to equal medoid distances. We resolve a medoid tie by input order and calculate centrality on the full QC-passing ensemble before support filtering. We keep these scores fixed during subset selection.

### B.2 Support (recurrence across folds)

We construct *B* = 20 reproducible partitions of the same ensemble into *K* = 5 disjoint folds. For each partition, we assign folds by a deterministic hash of the antibody identifier, sample identifier, and partition index. Let 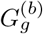 be fold *g* in partition *b*, and let 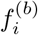 identify the fold containing member *i*. We find the nearest neighbor of *i* in each of the four other folds and take the median of these distances:

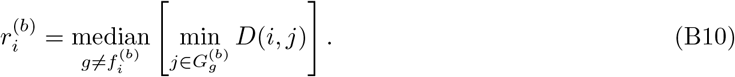

Small values indicate that the member has close conformational matches across several folds. The partitions reuse one generated ensemble; they do not correspond to separate generation runs.

To account for variation in neighbor spacing across the ensemble, we compare recurrence distances among members at similar distances from the medoid. We sort the members by *a*_*i*_ using a stable sort and define a local comparison window *W*_*i*_ around each member’s rank. The window contains *w*_0_ = min*{n*, max[51, *⌈*0.10*n⌉*]*}* members. We increase an even *w*_0_ by one when *w*_0_ *< n* and shift the window at either end of the ordering to preserve its size. Within each partition, we rank 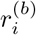 against the recurrence distances in *W*_*i*_, then take the median of these scores across partitions:

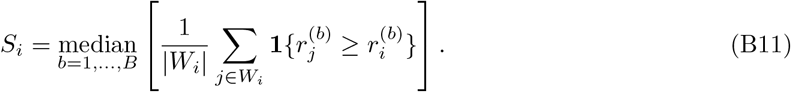

Higher support therefore indicates closer cross-fold recurrence than for other members at comparable medoid distances. This local comparison reduces the tendency to assign low support to peripheral members solely because their neighborhoods are sparser. It does not require support and centrality to be statistically independent.

## Appendix C Crystallography details

### C.1 Protein expression and purification, crystallization, and structure determination

Fab proteins were expressed and purified as previously described [61] and concentrated to 10 mg/mL, respectively, and crystallized in various PEG and ethylene glycol solutions. Crystals were cryocooled in liquid nitrogen, in their growth conditions supplemented with ethylene glycol. Diffraction data were collected at synchrotron sources, and those data were scaled using XDS [62] or HKL2000 [63]. The structures were determined by molecular replacement, using PHASER [64], and the models were adjusted in COOT [65] and refined in PHENIX [66] (Table C2). The data set for the Apo 1 structure showed significant anisotropy, with spherical completeness *>* 50% in the highest resolution bin occurring at 2.8 Å resolution and elliptically truncated data to 2.5 Å resolution utilized in refinement. The CDR regions of the Fabs were generally well ordered in the electron density maps (Figure C4).

**Table C2:**
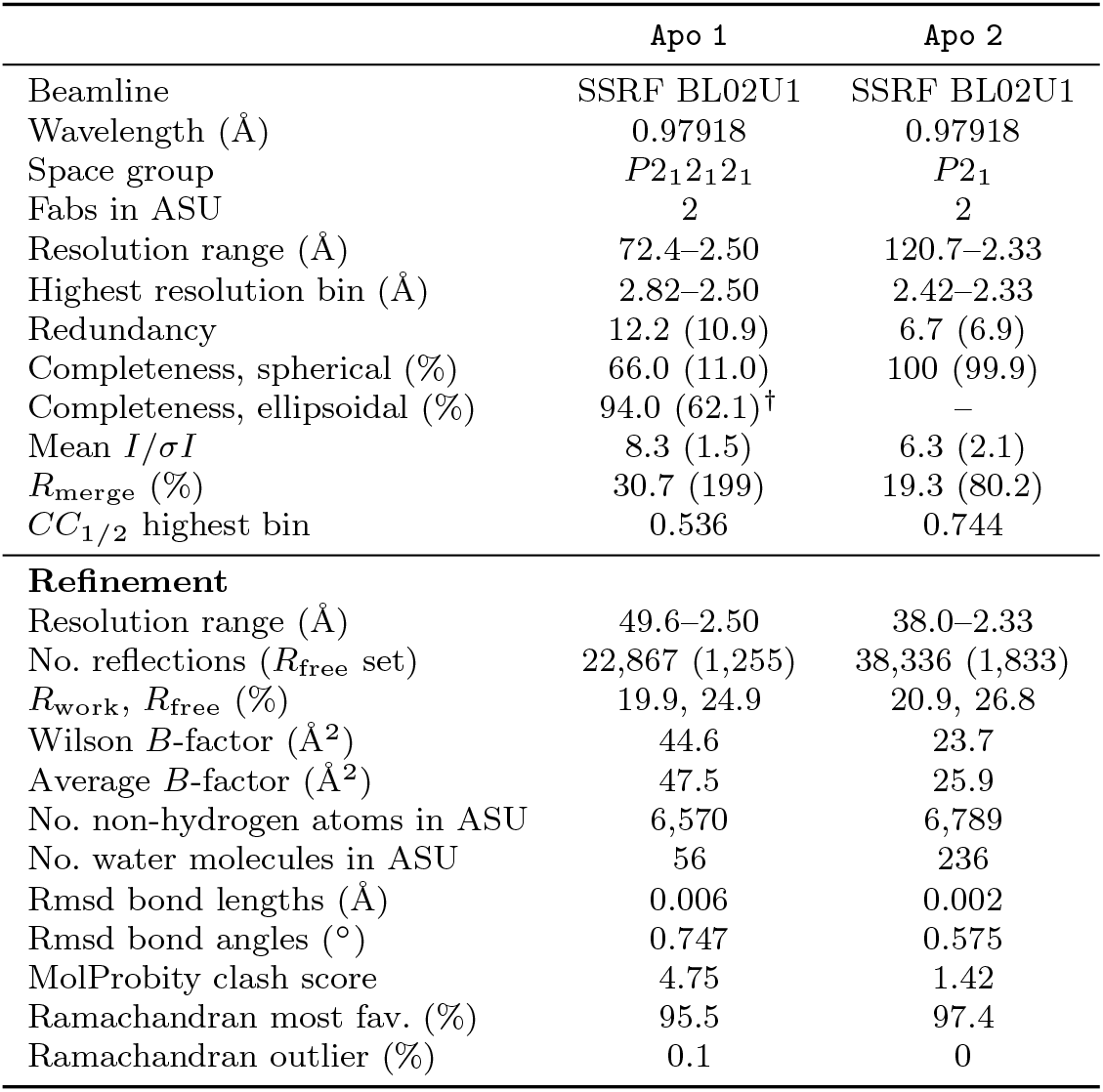
Crystallographic data and refinement statistics. Values in parentheses represent data from the highest resolution shell. ^*†*^Data were anisotropically scaled. Completeness reached *>* 50% in the highest resolution bin at 2.8 Å resolution.

|  | Apo 1 | Apo 2 |
|---|---|---|
| Beamline | SSRF BL02U1 | SSRF BL02U1 |
| Wavelength (Å) | 0.97918 | 0.97918 |
| Space group | *P*2<sub>1</sub>2<sub>1</sub>2<sub>1</sub> | *P*2<sub>1</sub> |
| Fabs in ASU | 2 | 2 |
| Resolution range (Å) | 72.4–2.50 | 120.7–2.33 |
| Highest resolution bin (Å) | 2.82–2.50 | 2.42–2.33 |
| Redundancy | 12.2 (10.9) | 6.7 (6.9) |
| Completeness, spherical (%) | 66.0 (11.0) | 100 (99.9) |
| Completeness, ellipsoidal (%) | 94.0 (62.1)<sup>†</sup> | – |
| Mean *I*/*σI* | 8.3 (1.5) | 6.3 (2.1) |
| *R*<sub>merge</sub> (%) | 30.7 (199) | 19.3 (80.2) |
| *CC*<sub>1/2</sub> highest bin | 0.536 | 0.744 |
| **Refinement** |  |  |
| Resolution range (Å) | 49.6–2.50 | 38.0–2.33 |
| No. reflections (*R*<sub>free</sub> set) | 22,867 (1,255) | 38,336 (1,833) |
| *R*<sub>work</sub>, *R*<sub>free</sub> (%) | 19.9, 24.9 | 20.9, 26.8 |
| Wilson *B*-factor (Å<sup>2</sup>) | 44.6 | 23.7 |
| Average *B*-factor (Å<sup>2</sup>) | 47.5 | 25.9 |
| No. non-hydrogen atoms in ASU | 6,570 | 6,789 |
| No. water molecules in ASU | 56 | 236 |
| Rmsd bond lengths (Å) | 0.006 | 0.002 |
| Rmsd bond angles (°) | 0.747 | 0.575 |
| MolProbity clash score | 4.75 | 1.42 |
| Ramachandran most fav. (%) | 95.5 | 97.4 |
| Ramachandran outlier (%) | 0.1 | 0 |

## Appendix D Large antibody molecular dynamics dataset

### D.1 Sequence selection

The dataset comprises 13,192 unique antibody Fv sequences drawn from three sources: the Structural Antibody Database (SAbDab) [52], the Therapeutic Structural Antibody Database (TheraSAbDab) [53], and the paired Observed Antibody Space (pOAS) [54]. From SAbDab and TheraSAbDab, we selected all non-redundant antibody sequences available as of January 2024. For pOAS, which contains over 2 million sequences, we applied two complementary subselection strategies. The first intentionally oversampled sequences with extreme surface patch properties, specifically Fv surface electrostatic potential (calculated via APBS) and spatial aggregation propensity (SAP_BM_) scores. The second maximized diversity across V-gene usage, Fv and CDR lengths, clonotype, and predicted biophysical properties including charge and hydrophobicity. All sequences were selected based on the Fv region.

### D.2 Structure prediction and refinement

Selected Fv sequences were folded using ESMFold (3B parameters) [31], which generates structure predictions end-to-end from sequence using the ESM-2 language model. To ensure Fv stability, sequences were folded as Fabs using the Herceptin constant domain as a template. Predicted structures were then minimized in Rosetta using the default score function with a custom disulfide bond energy weight of 10.0. Both backbone and sidechain dihedrals were allowed to move during minimization, which used the L-BFGS Armijo nonmonotone algorithm.

### D.3 Molecular dynamics simulations

Simulations were initiated from the ESMFold-predicted and Rosetta-minimized structures. Hydrogen atoms were added using PROPKA [67] at pH 7.4, and protonation states were assigned using PDB2PQR [68] with the Amber FF14SB force field [56]. Each antibody was solvated in a cubic box of TIP3P water with a 10 Å buffer from the protein surface, and the system was neutralized with NaCl ions.

**Fig. C4:**
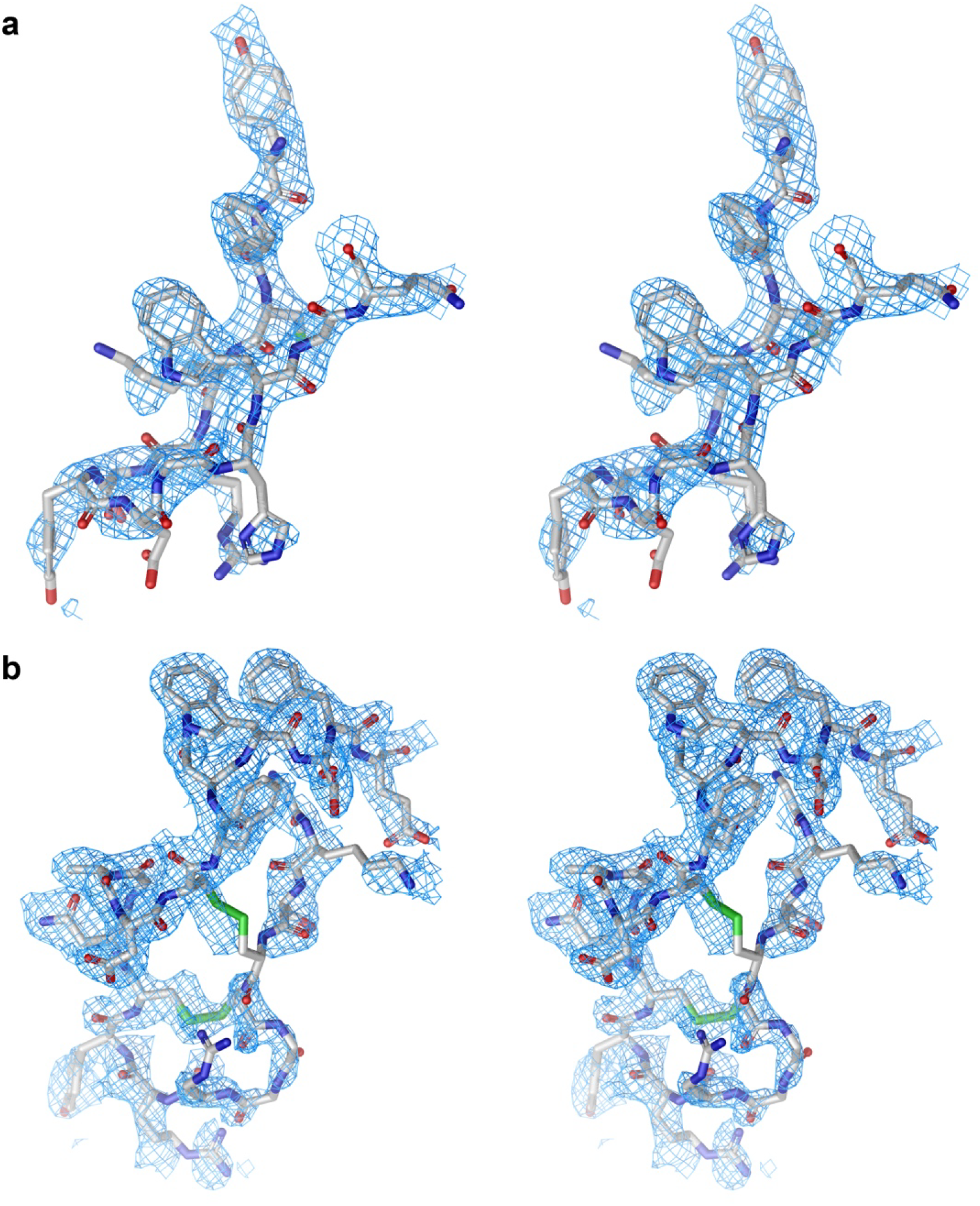
Divergent eye stereo diagram of the simulated annealing composite omit difference electron density maps, (2*m*|*F*_*o*_| *− D*|*F*_*c*_|) exp(*iα*_*c*_), of the CDR-H3 loops of **(a)** Apo 1 and **(b)** Apo 2.

System preparation proceeded as follows: (1) 2,000 steps of conjugate-gradient energy minimization; (2) 0.5 ns heating to 300 K under isothermal-isobaric (NPT) conditions with harmonic restraints on the protein (10 kcal mol^-1^ Å^-2^); (3) an additional 2,000 steps of unrestrained conjugate-gradient minimization; (4) 0.5 ns NPT equilibration at 1 atm and 300 K with restraints (10 kcal mol^-1^ Å^-2^); (5) 1 ns equilibration with reduced restraints (1 kcal mol^-1^ Å^-2^); and (6) final unrestrained equilibration at 300 K.

Production simulations were carried out under NPT conditions using hydrogen mass repartitioning (HMR), enabling a 4 fs timestep. Temperature was maintained at 298 K using Langevin dynamics with a collision frequency of 3 ps^-1^. Periodic boundary conditions were applied in all directions, electrostatic interactions were calculated using the particle mesh Ewald method [69] with a real-space cutoff of 10 Å, and hydrogen bond lengths were constrained with the SHAKE algorithm [70]. Each simulation was run for 200 ns on NVIDIA A100 GPUs, requiring approximately 12 hours per trajectory.

## Appendix E MSM and PCCA+ conformational clustering

We used a time-resolved clustering procedure to identify metastable conformational states in the molecular dynamics trajectories used for Cadense training. The models fitted here are used only to bias frame selection during training, not to estimate thermodynamic quantities. With 200 ns trajectories, the fitted MSMs give an approximate rather than a converged description of the underlying dynamics; we rely on them for the relative weighting of conformational states within a trajectory, and model quality is assessed against experimental structures and MD-derived observables rather than against the MSM itself.

For each trajectory, we extracted the backbone *ϕ* and *ψ* dihedral angles for the six antibody complementarity-determining regions (CDRs): H1, H2, H3, L1, L2, and L3. Because dihedral angles are periodic, we represented each angle *θ* using its sine and cosine,

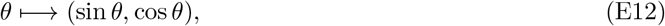

which avoids an artificial discontinuity between *π* and *π*. The resulting feature vector described the CDR backbone conformation at each trajectory frame.

### E.1 tICA projection and microstate assignment

We projected each dihedral feature trajectory onto time-lagged independent components using PyEMMA’s tICA implementation. We used a lag time of 50 frames and retained the first 30 components. The tICA coordinates were scaled using the kinetic-map convention, which weights each component according to its relaxation timescale and contribution to slow configurational dynamics. The trajectories used a frame spacing of 0.1 ns, so the tICA lag corresponded to 5 ns.

We then applied K-means clustering in the 30-dimensional tICA space. Each trajectory was partitioned into 150 microstates using a fixed random seed of zero. The number of microstates was fixed across all trajectories rather than selected per system. The resulting sequence of discrete microstate labels provided the input for Markov state model estimation. We used the tICA projection for both the dynamical state assignment and the subsequent selection of representative structures.

### E.2 Markov state model estimation

We estimated a reversible Markov state model for each trajectory from its discrete microstate sequence; reversibility is imposed by the estimator rather than tested. To select the MSM lag time, we evaluated lag times between 0.1 and 2.0 ns. For each candidate lag time, we calculated the leading five implied timescales, and selected the first lag time at which all five changed by less than 20% relative to the next tested lag time. The selected lag time could therefore vary between trajectories. We treat this as a practical heuristic for lag-time selection rather than as evidence of Markovianity; at these trajectory lengths the implied timescales are not expected to be fully converged.

At the selected lag time, we fit a reversible Bayesian Markov state model using 200 posterior samples. When Bayesian estimation failed, we used the corresponding reversible maximum-likelihood MSM. The stationary distribution of the fitted MSM provided a stationary weight for each microstate under that model.

### E.3 PCCA+ macrostates and representative structures

We applied PCCA+ to each fitted MSM and grouped the 150 microstates into 12 metastable macrostates, again using a fixed number across all trajectories. PCCA+ uses the slow dynamical modes of the MSM to group microstates that exchange rapidly with one another while separating states connected by slower transitions. For each macrostate *M*_*k*_, we calculated its stationary weight by summing the stationary probabilities of its constituent microstates,

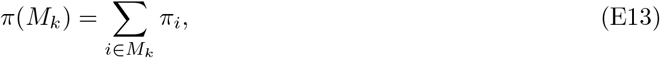

where *π*_*i*_ denotes the MSM stationary probability of microstate *i*. We normalized the resulting macrostate weights for use during training-frame selection.

We selected up to 10 structural representatives from each macrostate. For every microstate, we identified the trajectory frame closest to its K-means centroid in tICA space. We calculated the centroid of the microstates assigned to each PCCA+ macrostate and selected the microstate representatives closest to this macrostate centroid. The corresponding trajectory frames were written as representative PDB structures. This procedure preserved the dynamical organization supplied by the MSM and selected structures near the geometric center of each metastable state.

The resulting PCCA+ macrostate weights defined the curriculum used for the primary Cadense model. Frames associated with more heavily weighted macrostates received greater sampling weight, whereas uniformly spaced frame selection was used for the Cadense-*dist* variant. The two models therefore used the same structural and dihedral representations but differed in whether MD frames were sampled according to MSM-derived stationary weights or temporal spacing.

## Appendix F Model architecture details

### F.1 Protein geometry parameterization

We parameterize each residue by a backbone rigid frame—a translation *t*_*i*_ *∈* ℝ^3^ and rotation *r*_*i*_ *∈* SO(3), derived from the C_*α*_, N, and C backbone heavy atoms following AlphaFold2 Algorithm 21 [4]—together with five torsion angles *χ*_*iq*_ *∈* SO(2) for *q ∈ {*0, …, 4*}*. Here *χ*_0_ *≡ ψ* is the backbone oxygen torsion and *χ*_1_, …, *χ*_4_ are the four side-chain dihedral angles. For residue types with fewer than four side-chain torsions, the extra angles are masked during inference according to the fixed residue identity. The state of a system is specified by *{t*_*i*_*}, {r*_*i*_*}, {χ*_*iq*_*}*, collectively denoted *Z*.

### F.2 Input feature embedding

The `InputFeatureEmbedder` initializes single (per-residue, 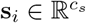) and pair (per-residue-pair, 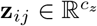) representations for the input conditioner. Single representations are constructed from:

- One-hot encoded amino acid identity
- Protein language model embeddings from ESM-2 [31]
- Per-residue confidence features
- Noise level embeddings via learned lookup tables

These are concatenated and projected through a linear layer (without bias). Pair representations are initialized from relative position encodings based on residue index and chain index, following AlphaFold2 and AlphaFold3 [4, 10].

### F.3 Input conditioner

The input conditioner constitutes the computational bulk of the model. Single and pair embeddings from the `InputFeatureEmbedder` are passed through a 1D–2D mixing layer (outer product of single representations added to pair representations), then refined through *N*_cycles_ Pairformer blocks. Each Pairformer block includes triangle attention across rows and columns of the pair representation, triangle multiplicative updates, and single-representation attention, analogous to the Evoformer/Pairformer stacks in AlphaFold2/3 [4, 10].

To improve computational efficiency, we use semi-local variants of triangle attention and triangle multiplication following Proteus [43]. Attention coefficients, which are rank-3 tensors for triangle attention, are sparsified using a neighborhood sampling scheme: *k*-nearest sequence neighbors plus random graph edges (without radial propensity sampling, to avoid leaking geometric information into the input conditioner). This reduces the computational complexity from *O* (*n*^3^) to *O* (*kn*^2^) for coordination number *k* ≤ *n* and *n* residues.

At each recycling iteration, the single and pair representations from the previous cycle are added (after layer normalization and linear projection) to the current representations before entering the Pairformer stack. The final single *{***s**_*i*_*}* and pair *{***z**_*ij*_*}* embeddings, along with their initial representations 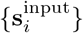 and 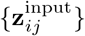, serve as conditioning information for the diffusion module.

### F.4 Diffusion conditioner

At each diffusion timestep, the conditioning embeddings are updated by the `DiffusionConditioner` before being passed to the score model. This module:

1. Concatenates current single/pair embeddings with their initial (pre-Pairformer) versions
2. Adds self-conditioning information: the output single and pair embeddings from the previous diffusion timestep (initialized to zero for the first step), following RFdiffusion [71] and Proteus [43]
3. Incorporates a learned time embedding via sinusoidal encoding projected through layer normalization and a linear layer
4. Refines through *N*_transition_ transition blocks (2-layer MLPs with residual connections)

### F.5 Score model

The score model predicts denoising scores for all geometric degrees of freedom given the current noised structure *Z*^(*t*)^ and conditioning embeddings.

#### Graph construction

Backbone coordinates *t*_*i*_ are used to construct a message-passing graph *G* with *k* = 32 nearest-neighbor edges plus *f* = 32 random long-range edges sampled with inverse-cubic radial propensity, following Chroma [45].

#### Geometric featurization

The pair embeddings **z**_*ij*_ within the score model are augmented with geometric features derived from the current structure, including backbone distance (distogram), planar angles, and dihedral angles between connected residues. These provide the score model with information about the current structural state at each diffusion timestep, biasing the local triangle attention updates with physically relevant geometric context.

#### Local Pairformer

The conditioning single and pair embeddings are first refined through Pairformer blocks with triangle attention restricted to the edges of the local graph *G*, reducing computational cost relative to full-attention Pairformer blocks in the input conditioner.

#### Equivariant layers

The core of the score model uses the Equiformer architecture [44], a graph attention network with SE(3)-equivariant message passing based on irreducible representations of SO(3). We restrict steerable features to maximum order *L*_max_ = 1, i.e., scalar (*L* = 0) and vector (*L* = 1) features, and use depth-wise Clebsch– Gordan tensor products to reduce computational complexity. (We do not use the eSCN tensor product simplifications from Equiformer V2 [72], as the gains are minimal at *L*_max_ = 1.)

#### Global frame construction

Side-chain torsion angles are mapped to global SE(3) frames following Equifold [46], producing 6 frames per residue (1 backbone + up to 5 torsion-derived). Each global frame is assigned a node in the equivariant layers with initial features derived from the backbone rotation and sequence embeddings.

#### Hierarchical message passing

We employ three tiers of nodes in the equivariant layers:

1. **Backbone (BB) nodes**: one per residue, connected via the Chroma-style graph *G*
2. **Side-chain (SC) nodes**: one per global frame (excluding the backbone frame), connected *only* to their parent BB node. Initial BB featurization includes a directed message-passing step from SC to BB nodes
3. **Global auxiliary node**: a single node holding a latent representation of the full system state, fully connected to all BB nodes

Message passing alternates between: (1) directed aggregation of all BB features to the auxiliary node, and (2) standard message passing across the combined graph of BB nodes and the auxiliary node. This hierarchy avoids the *O* (*n*_BB_ *×n*_SC_) cost that would arise from building a full radial graph over all backbone and side-chain nodes, while still enabling side-chain information to influence backbone denoising through the hierarchical update.

## Appendix G Diffusion process details

### G.1 Forward processes and noise schedules

We apply independent diffusion processes to each type of geometric parameter, using a variance-preserving (VP) process for translations and variance-exploding (VE) processes for rotations and torsions, following Song et al. [50].

#### Translations (VP)

Backbone translations *t*_*i*_ are constrained to zero center-of-mass (∑_*i*_ *t*_*i*_ = 0), following FrameDiff [49]. The noise schedule follows a geometric progression with an implicit physical length scale, parameterized following Kingma et al. [48]. The stationary distribution is a zero-center-of-mass projected normal with isotropic diagonal covariance scaled by physical units. Minimum and maximum noise scales are 0.05 Å and 300.0 Å, respectively.

#### Rotations (VE)

Backbone rotations follow Brownian motion on SO(3) with a variance-exploding schedule [50]. The stationary distribution is the uniform Haar measure, i.e., the high-noise limit of the IGSO(3) heat kernel. Minimum noise scale is 0.035 rad.

#### Torsions (VE)

Side-chain and backbone oxygen torsion angles follow Brownian motion on SO(2) with a VE schedule [50], following DiffDock [51]. The stationary distribution is the uniform measure on SO(2) (the high-noise limit of the wrapped normal distribution). Torsion angles with *π*-periodicity are handled specially in loss calculations based on residue identity. Minimum noise scale is 0.035 rad.

#### Schedule calibration

Setting noise schedules simultaneously across parameter types is nontrivial. Following Biroli et al. [73], we calibrated schedules so that the symmetry-breaking phase transitions for different degrees of freedom occur at approximately the same diffusion time. We estimated approximate transition scales as 1.0 Å for translations and 0.2 rad for rotations and torsions. Given minimum noise levels (set to expected data noise scales) and these transition scales, maximum noise levels were determined from the functional forms of the schedules and further adjusted empirically.

### G.2 Correlated noise for zero-center-of-mass translations

The zero-center-of-mass constraint on translations induces correlations in the noise distribution. The covariance matrix of the stationary distribution for translations within a residue group of *n* residues is given by 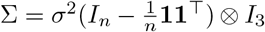, where *σ* is the schedule-dependent noise scale. Sampling is performed by drawing from an unconstrained normal and projecting onto the zero-center-of-mass subspace.

## Appendix H Sampling details

### H.1 Reverse diffusion

Baseline reverse-diffusion sampling uses Euler–Maruyama discretization of the time-reversed SDEs over SE(3) *×* SO(2)^5^. We use evenly spaced timesteps (corresponding to *ρ* = 1 in the EDM parameterization of Karras et al. [74]).

### H.2 Hybrid Langevin dynamics

Following Chroma [45], we implement hybrid Langevin dynamics to control the diversity–quality tradeoff. The target distribution is rescaled as 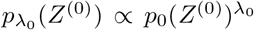, where *λ*_0_ *>* 1 is a low-temperature regime emphasizing high-likelihood states. This induces a time-dependent rescaling of the scores of the form *λ*_*t*_ = *F* (*λ*_0_, *σ*_*t*_), where the functional form of *F* depends on the noise schedule and the manifold of the given degrees of freedom.

To compensate for inadequate reweighting of relative populations across modes under simple score rescaling, we introduce an equilibration factor *ψ >* 0 that adds a component of slow, annealed Langevin dynamics [50]. The reverse step under hybrid dynamics takes the form:

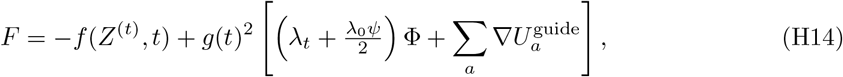

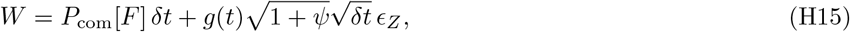

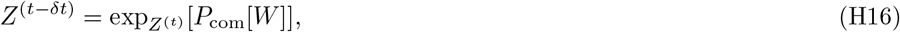

where Φ denotes the predicted scores, *P*_com_ is the zero-center-of-mass projection, *ϵ*_*Z*_ is noise sampled on the appropriate manifold, and exp is the Riemannian exponential map.

Corrector steps follow annealed Langevin dynamics at fixed noise level *t*, using the same form but without the drift term *f* and with unit noise scaling.

### H.3 Guidance potentials

At inference time, auxiliary differentiable potentials 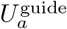 can be composed with the learned score to guide sampling. Gradients 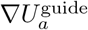 are added to the score in both predictor and corrector steps. We use guidance based on the AlphaFold2 structure violation loss [4], which penalizes steric clashes and bond geometry violations, to improve the physical quality of generated structures. When guidance is active, we found it necessary to include corrector steps (up to 5 per predictor step) to allow the reverse dynamics to mix adequately under the additional gradient field.

## Footnotes

1 We found Ibex-*holo* outperformed the *apo* version for docking, consistent with [8], so we omit the latter in this section.

## References

[1] Carter, P.J., Rajpal, A.: Designing antibodies as therapeutics. Cell 185(15), 2789–2805 (2022)

[2] Sharma, P., Joshi, R.V., Pritchard, R., Xu, K., Eicher, M.A.: Therapeutic antibodies in medicine. Molecules 28(18), 6438 (2023)

[3] Nelson, A.L., Dhimolea, E., Reichert, J.M.: Development trends for human monoclonal antibody therapeutics. Nature reviews drug discovery 9(10), 767–774 (2010)

[4] Jumper, J., Evans, R., Pritzel, A., Green, T., Figurnov, M., Ronneberger, O., Tunyasuvunakool, K., Bates, R., Žídek, A., Potapenko, A., et al.: Highly accurate protein structure prediction with alphafold. nature 596(7873), 583–589 (2021)

[5] Abanades, B., Wong, W.K., Boyles, F., Georges, G., Bujotzek, A., Deane, C.M.: Immunebuilder: Deep-learning models for predicting the structures of immune proteins. Communications Biology 6(1), 575 (2023)

[6] Kenlay, H., Dreyer, F.A., Cutting, D., Nissley, D., Deane, C.M.: Abodybuilder3: improved and scalable antibody structure predictions. Bioinformatics 40(10), 576 (2024)

[7] Ruffolo, J.A., Chu, L.-S., Mahajan, S.P., Gray, J.J.: Fast, accurate antibody structure prediction from deep learning on massive set of natural antibodies. Nature communications 14(1), 2389 (2023)

[8] Dreyer, F.A., Ludwiczak, J., Martinkus, K., Abanades, B., Alberstein, R.G., Kessel, P., Rao, P., Lee, J.H., Bonneau, R., Watkins, A.M., et al.: Conformation-aware structure prediction of antigen-recognizing immune proteins. In: Mabs, vol. 18, p. 2602217 (2026). Taylor & Francis

[9] Hummer, A.M., Abanades, B., Deane, C.M.: Advances in computational structure-based antibody design. Current opinion in structural biology 74, 102379 (2022)

[10] Abramson, J., Adler, J., Dunger, J., Evans, R., Green, T., Pritzel, A., Ronneberger, O., Will-more, L., Ballard, A.J., Bambrick, J., Bodenstein, S.W., Evans, D.A., Hung, C.-C., O’Neill, M., Reiman, D., Tunyasuvunakool, K., Wu, Z., Žemgulytė, A., Arvaniti, E., Beattie, C., Bertolli, O., Bridgland, A., Cherepanov, A., Congreve, M., Cowen-Rivers, A.I., Cowie, A., Figurnov, M., Fuchs, F.B., Gladman, H., Jain, R., Khan, Y.A., Low, C.M.R., Perlin, K., Potapenko, A., Savy, P., Singh, S., Stecula, A., Thillaisundaram, A., Tong, C., Yakneen, S., Zhong, E.D., Zielinski, M., Žídek, A., Bapst, V., Kohli, P., Jaderberg, M., Hassabis, D., Jumper, J.M.: Accurate structure prediction of biomolecular interactions with alphafold 3. Nature 630(8016), 493–500 (2024) 10.1038/s41586-024-07487-w

[11] Wohlwend, J., Corso, G., Passaro, S., Reveiz, M., Leidal, K., Swiderski, W., Portnoi, T., Chinn, I., Silterra, J., Jaakkola, T., Barzilay, R.: Boltz-1 democratizing biomolecular interaction modeling. bioRxiv (2024) 10.1101/2024.11.19.624167 https://www.biorxiv.org/content/early/2024/11/20/2024.11.19.624167.full.pdf

[12] team, C.D., Boitreaud, J., Dent, J., McPartlon, M., Meier, J., Reis, V., Rogozhonikov, A., Wu K.: Chai-1: Decoding the molecular interactions of life. bioRxiv (2024) 10.1101/2024.10.10.615955 https://www.biorxiv.org/content/early/2024/10/11/2024.10.10.615955.full.pdf

[13] Protenix Team, Zhang, Y., Gong, C., Zhang, H., Ma, W., Liu, Z., Chen, X., Guan, J., Wang, L., Xiao, W.: Protenix-v1: Toward high-accuracy open-source biomolecular structure prediction. bioRxiv (2026) 10.64898/2026.02.05.703733

[14] Team, C.D., Boitreaud, J., Dent, J., Geisz, D., McPartlon, M., Meier, J., Qiao, Z., Rogozhnikov, A., Rollins, N., Wollenhaupt, P., et al.: Zero-shot antibody design in a 24-well plate. bioRxiv, 2025–07 (2025)

[15] Mille-Fragoso, L.S., Wang, J.N., Driscoll, C.L., Dai, H., Widatalla, T., Zhang, X., Hie, B.L., Gao, X.J.: Efficient generation of epitope-targeted de novo antibodies with germinal. bioRxiv (2025)

[16] Li, T., Tracka, M.B., Uddin, S., Casas-Finet, J., Jacobs, D.J., Livesay, D.R.: Rigidity emerges during antibody evolution in three distinct antibody systems: evidence from qsfr analysis of fab fragments. PLoS computational biology 11(7), 1004327 (2015)

[17] Fernández-Quintero, M.L., Loeffler, J.R., Bacher, L.M., Waibl, F., Seidler, C.A., Liedl, K.R.: Local and global rigidification upon antibody affinity maturation. Frontiers in molecular biosciences 7, 182 (2020)

[18] Blackler, R.J., Müller-Loennies, S., Pokorny-Lehrer, B., Legg, M.S., Brade, L., Brade, H., Kosma, P., Evans, S.V.: Antigen binding by conformational selection in near-germline antibodies. Journal of Biological Chemistry 298(5), 101901 (2022)

[19] Spoendlin, F.C., Fernández-Quintero, M.L., Raghavan, S.S., Turner, H.L., Gharpure, A., Loeffler, J.R., Wong, W.K., Bujotzek, A., Georges, G., Ward, A.B., et al.: Predicting the conformational flexibility of antibody and t-cell receptor cdrs. bioRxiv, 2025–03 (2025)

[20] Cagiada, M., Spoendlin, F.C., Ifashe, K., Deane, C.M.: Uncovering the flexibility of cdr loops in antibodies and tcrs through large-scale molecular dynamics. bioRxiv, 2025–11 (2025)

[21] Fernandez-Quintero, M.L., Vangone, A., Loeffler, J.R., Seidler, C.A., Georges, G., Liedl, K.R.: Paratope states in solution improve structure prediction and docking. Structure 30(3), 430–440 (2022)

[22] Park, E., Izadi, S.: Molecular surface descriptors to predict antibody developability: sensitivity to parameters, structure models, and conformational sampling. In: MAbs, vol. 16, p. 2362788 (2024). Taylor & Francis

[23] Wang, A., Sang, Z., Stanton, S.D., Hofmann, J.L., Izadi, S., Park, E., Ludwiczak, J., Kirchmeyer, M., Davidson, D., Maier, A., et al.: A guided design framework for the optimization of therapeutic-like antibodies. In: ICLR 2025 Workshop on Generative and Experimental Perspectives for Biomolecular Design (2025)

[24] Rao, P., Isaacson, H., Hofmann, J.L., Davidson, D., Wang, A., Watkins, A.M., Bonneau, R., Izadi, S., Lee, J.H.: Surfprop: A surface-based property prediction framework for antibody developability and screening. In: ICML 2025 Generative AI and Biology (GenBio) Workshop (2025)

[25] Jing, B., Berger, B., Jaakkola, T.: Alphafold meets flow matching for generating protein ensembles. In: Forty-first International Conference on Machine Learning (2024)

[26] Lewis, S., Hempel, T., Jiménez-Luna, J., Gastegger, M., Xie, Y., Foong, A.Y., Satorras, V.G., Abdin, O., Veeling, B.S., Zaporozhets, I., et al.: Scalable emulation of protein equilibrium ensembles with generative deep learning. Science 389(6761), 9817 (2025)

[27] Passaro, S., Corso, G., Wohlwend, J., Reveiz, M., Thaler, S., Ram Somnath, V., Getz, N., Portnoi, T., Roy, J., Stark, H., et al.: Boltz-2: Towards accurate and efficient binding affinity prediction. bioRxiv, 2025–06 (2025)

[28] Spoendlin, F.C., Cagiada, M., Ifashe, K., Vavourakis, O., Deane, C.M.: Sampling antibody conformational ensembles with abodybuilder4-steroids. bioRxiv, 2026–04 (2026)

[29] Xu, X., Giulini, M., Bonvin, A.M.: Improved prediction of antibody and their complexes with clustered generative modelling ensembles. Bioinformatics Advances 5(1), 161 (2025)

[30] Borges-Araújo, L., Patmanidis, I., Singh, A.P., Santos, L.H., Sieradzan, A.K., Vanni, S., Czaplewski, C., Pantano, S., Shinoda, W., Monticelli, L., et al.: Pragmatic coarse-graining of proteins: models and applications. Journal of Chemical Theory and Computation 19(20), 7112–7135 (2023)

[31] Lin, Z., Akin, H., Rao, R., Hie, B., Zhu, Z., Lu, W., Smetanin, N., Verkuil, R., Kabeli, O., Shmueli, Y., et al.: Evolutionary-scale prediction of atomic-level protein structure with a language model. Science 379(6637), 1123–1130 (2023)

[32] Richman, D.D., Karaguesian, J., Suomivuori, C.-M., Dror, R.O.: Unlocking hidden biomolecular conformational landscapes in diffusion models at inference time. arXiv preprint arXiv:2512.03312 (2025)

[33] Hitawala, F.N., Gray, J.J.: What does alphafold3 learn about antigen and nanobody docking, and what remains unsolved? bioRxiv, 2024–09 (2025)

[34] Kelow, S., Faezov, B., Xu, Q., Parker, M., Adolf-Bryfogle, J., Dunbrack Jr, R.L.: A penultimate classification of canonical antibody cdr conformations. biorxiv, 2022–10 (2022)

[35] North, B., Lehmann, A., Dunbrack Jr, R.L.: A new clustering of antibody cdr loop conformations. Journal of molecular biology 406(2), 228–256 (2011)

[36] Giulini, M., Reys, V., Teixeira, J.M., Jiménez-García, B., V. Honorato, R., Kravchenko, A., Xu, X., Versini, R., Engel, A., Verhoeven, S., et al.: Haddock3: A modular and versatile platform for integrative modeling of biomolecular complexes. Journal of Chemical Information and Modeling (2025)

[37] Basu, S., Wallner, B.: Dockq: a quality measure for protein-protein docking models. PloS one 11(8), 0161879 (2016)

[38] Xu, X., Coratella, I., Bonvin, A.M.: Deeprank-ab: a dedicated scoring function for antibody-antigen complexes based on geometric deep learning. bioRxiv, 2025–12 (2025)

[39] Tadiello, M., Ludaic, M., Viliuga, V., Elofsson, A.: Abag-rank: Improving model selection of alphafold antibody–antigen complexes by learning to rank. bioRxiv, 2026–03 (2026)

[40] Ishima, R., Torchia, D.A.: Protein dynamics from nmr. Nature structural biology 7(9), 740–743 (2000)

[41] Walters, B.T., Patapoff, A.W., Kiefer, J.R., Wu, P., Wang, W.: Integrating hydrogen exchange with molecular dynamics for improved ligand binding predictions. Journal of Chemical Information and Modeling (2025)

[42] Liu, Y., Yu, Z., Lindsay, R.J., Lin, G., Chen, M., Sahoo, A., Hanson, S.M.: Exendiff: an experiment-guided diffusion model for protein conformational ensemble generation. PRX Life 3(2), 023013 (2025)

[43] Wang, C., Qu, Y., Peng, Z., Wang, Y., Zhu, H., Chen, D., Cao, L.: Proteus: exploring protein structure generation for enhanced designability and efficiency. bioRxiv, 2024–02 (2024)

[44] Liao, Y.-L., Smidt, T.: Equiformer: Equivariant graph attention transformer for 3d atomistic graphs. arXiv preprint arXiv:2206.11990 (2022)

[45] Ingraham, J.B., Baranov, M., Costello, Z., Barber, K.W., Wang, W., Ismail, A., Frappier, V., Lord, D.M., Ng-Thow-Hing, C., Van Vlack, E.R., et al.: Illuminating protein space with a programmable generative model. Nature 623(7989), 1070–1078 (2023)

[46] Lee, J.H., Yadollahpour, P., Watkins, A., Frey, N.C., Leaver-Fay, A., Ra, S., Cho, K., Gligorijević, V., Regev, A., Bonneau, R.: Equifold: Protein structure prediction with a novel coarse-grained structure representation. Biorxiv, 2022–10 (2022)

[47] De Bortoli, V., Mathieu, E., Hutchinson, M., Thornton, J., Teh, Y.W., Doucet, A.: Riemannian score-based generative modelling. Advances in neural information processing systems 35, 2406–2422 (2022)

[48] Kingma, D., Salimans, T., Poole, B., Ho, J.: Variational diffusion models. Advances in neural information processing systems 34, 21696–21707 (2021)

[49] Yim, J., Trippe, B.L., De Bortoli, V., Mathieu, E., Doucet, A., Barzilay, R., Jaakkola, T.: Se (3) diffusion model with application to protein backbone generation. arXiv preprint arXiv:2302.02277 (2023)

[50] Song, Y., Sohl-Dickstein, J., Kingma, D.P., Kumar, A., Ermon, S., Poole, B.: Score-based generative modeling through stochastic differential equations. arXiv preprint arXiv:2011.13456 (2020)

[51] Corso, G., Stärk, H., Jing, B., Barzilay, R., Jaakkola, T.: Diffdock: Diffusion steps, twists, and turns for molecular docking. arXiv preprint arXiv:2210.01776 (2022)

[52] Dunbar, J., Krawczyk, K., Leem, J., Baker, T., Fuchs, A., Georges, G., Shi, J., Deane, C.M.: Sabdab: the structural antibody database. Nucleic acids research 42(D1), 1140–1146 (2014)

[53] Raybould, M.I., Marks, C., Lewis, A.P., Shi, J., Bujotzek, A., Taddese, B., Deane, C.M.: Thera-sabdab: the therapeutic structural antibody database. Nucleic acids research 48(D1), 383–388 (2020)

[54] Olsen, T.H., Boyles, F., Deane, C.M.: Observed antibody space: A diverse database of cleaned, annotated, and translated unpaired and paired antibody sequences. Protein Science 31(1), 141–146 (2022)

[55] Leman, J.K., Weitzner, B.D., Lewis, S.M., Adolf-Bryfogle, J., Alam, N., Alford, R.F., Aprahamian, M., Baker, D., Barlow, K.A., Barth, P., et al.: Macromolecular modeling and design in rosetta: recent methods and frameworks. Nature methods 17(7), 665–680 (2020)

[56] Maier, J.A., Martinez, C., Kasavajhala, K., Wickstrom, L., Hauser, K.E., Simmerling, C.: ff14sb: improving the accuracy of protein side chain and backbone parameters from ff99sb. Journal of chemical theory and computation 11(8), 3696–3713 (2015)

[57] Pérez-Hernández, G., Paul, F., Giorgino, T., De Fabritiis, G., Noé, F.: Identification of slow molecular order parameters for markov model construction. The Journal of chemical physics 139(1) (2013)

[58] Deuflhard, P., Weber, M.: Robust perron cluster analysis in conformation dynamics. Linear algebra and its applications 398, 161–184 (2005)

[59] Scherer, M.K., Trendelkamp-Schroer, B., Paul, F., Pérez-Hernández, G., Hoffmann, M., Plattner, N., Wehmeyer, C., Prinz, J.-H., Noé, F.: PyEMMA 2: A Software Package for Estimation, Validation, and Analysis of Markov Models. Journal of Chemical Theory and Computation 11, 5525–5542 (2015) 10.1021/acs.jctc.5b00743. Accessed 2015-10-19

[60] Steinegger, M., Söding, J.: Mmseqs2 enables sensitive protein sequence searching for the analysis of massive data sets. Nature biotechnology 35(11), 1026–1028 (2017)

[61] Lin, J.Y.-Y., Hofmann, J.L., Leaver-Fay, A., Liang, W.-C., Vasilaki, S., Lee, E., Pinheiro, P.O., Tagasovska, N., Kiefer, J.R., Wu, Y., Seeger, F., Bonneau, R., Gligorijevic, V., Watkins, A.M., Cho, K., Frey, N.C.: DyAb: sequence-based antibody design and property prediction in a low-data regime. bioRxiv (2025) 10.1101/2025.01.28.635353.2025.01.28.635353

[62] Kabsch, W.: XDS. Acta Crystallographica Section D: Biological Crystallography 66, 125–132 (2010)

[63] Otwinowski, Z., Minor, W.: Processing of x-ray diffraction data collected in oscillation mode. Methods in Enzymology 276, 307–326 (1997)

[64] McCoy, A.J., Oeffner, R.D., Wrobel, A.G., Ojala, J.R.M., Tryggvason, K., Lohkamp, B., Read, R.J.: Ab initio solution of macromolecular crystal structures without direct methods. Proceedings of the National Academy of Sciences USA 114, 3637–3641 (2017)

[65] Emsley, P., Cowtan, K.: Coot: model-building tools for molecular graphics. Acta Crystallographica Section D: Biological Crystallography 60, 2126–2132 (2004)

[66] Adams, P.D., Afonine, P.V., Bunkóczi, G., Chen, V.B., Davis, I.W., Echols, N., Headd, J.J., Hung, L.-W., Kapral, G.J., Grosse-Kunstleve, R.W., McCoy, A.J., Moriarty, N.W., Oeffner, R., Read, R.J., Richardson, D.C., Richardson, J.S., Terwilliger, T.C., Zwart, P.H.: PHENIX: a comprehensive python-based system for macromolecular structure solution. Acta Crystallographica Section D: Biological Crystallography 66, 213–221 (2010)

[67] Olsson, M.H., Søndergaard, C.R., Rostkowski, M., Jensen, J.H.: Propka3: consistent treatment of internal and surface residues in empirical p k a predictions. Journal of chemical theory and computation 7(2), 525–537 (2011)

[68] Dolinsky, T.J., Nielsen, J.E., McCammon, J.A., Baker, N.A.: Pdb2pqr: an automated pipeline for the setup of poisson–boltzmann electrostatics calculations. Nucleic acids research 32(suppl 2), 665–667 (2004)

[69] Petersen, H.G.: Accuracy and efficiency of the particle mesh ewald method. The Journal of chemical physics 103(9), 3668–3679 (1995)

[70] Forester, T.R., Smith, W.: Shake, rattle, and roll: efficient constraint algorithms for linked rigid bodies. Journal of computational chemistry 19(1), 102–111 (1998)

[71] Watson, J.L., Juergens, D., Bennett, N.R., Trippe, B.L., Yim, J., Eisenach, H.E., Ahern, W., Borst, A.J., Ragotte, R.J., Milles, L.F., et al.: De novo design of protein structure and function with rfdiffusion. Nature 620(7976), 1089–1100 (2023)

[72] Liao, Y.-L., Wood, B., Das, A., Smidt, T.: Equiformerv2: Improved equivariant transformer for scaling to higher-degree representations. arXiv preprint arXiv:2306.12059 (2023)

[73] Biroli, G., Bonnaire, T., De Bortoli, V., Mézard, M.: Dynamical regimes of diffusion models. Nature Communications 15(1), 9957 (2024)

[74] Karras, T., Aittala, M., Aila, T., Laine, S.: Elucidating the design space of diffusion-based generative models. Advances in neural information processing systems 35, 26565–26577 (2022)

